# Mapping the genetic landscape establishing a tumor immune microenvironment favorable for anti-PD-1 response in mice and humans

**DOI:** 10.1101/2024.07.11.603136

**Authors:** Daniel A. Skelly, John P. Graham, Mingshan Cheng, Mayuko Furuta, Andrew Walter, Thomas A. Stoklasek, Hongyuan Yang, Timothy M. Stearns, Olivier Poirion, Ji-Gang Zhang, Jessica D. S. Grassmann, Diane Luo, William F. Flynn, Elise T. Courtois, Chih-Hao Chang, David V. Serreze, Francesca Menghi, Laura G. Reinholdt, Edison T. Liu

**Affiliations:** The Jackson Laboratory for Mammalian Genetics, Bar Harbor, ME, USA; The Jackson Laboratory, Sacramento, CA, USA; The Jackson Laboratory for Genomic Medicine, Farmington, CT, USA; Single Cell Biology Lab, The Jackson Laboratory for Genomic Medicine, Farmington, CT, USA; OB/Gyn Department, UConn Health, Farmington, CT, USA

## Abstract

Identifying host genetic factors modulating immune checkpoint inhibitor (ICI) efficacy has been experimentally challenging because of variations in both host and tumor genomes, differences in the microbiome, and patient life exposures. Utilizing the Collaborative Cross (CC) multi-parent mouse genetic resource population, we developed an approach that fixes the tumor genomic configuration while varying host genetics. With this approach, we discovered that response to anti-PD-1 (aPD1) immunotherapy was significantly heritable in four distinct murine tumor models (*H*^2^ between 0.18-0.40). For the MC38 colorectal carcinoma system (*H*^2^ = 0.40), we mapped four significant ICI response quantitative trait loci (QTL) localized to mouse chromosomes (mChr) 5, 9, 15 and 17, and identified significant epistatic interactions between specific QTL pairs. Differentially expressed genes within these QTL were highly enriched for immune genes and pathways mediating allograft rejection and graft vs host disease. Using a cross species analytical approach, we found a core network of 48 genes within the four QTLs that showed significant prognostic value for overall survival in aPD1 treated human cohorts that outperformed all other existing validated immunotherapy biomarkers, especially in human tumors of the previously defined immune subtype 4. Functional blockade of two top candidate immune targets within the 48 gene network, GM-CSF and high affinity IL-2/IL-15 signaling, completely abrogated the MC38 tumor transcriptional response to aPD1 therapy *in vivo*. Thus, we have established a powerful cross species *in vivo* platform capable of uncovering host genetic factors that establish the tumor immune microenvironment configuration propitious for ICI response.

## Introduction

Abrogating immune checkpoint control has proven to be a powerful therapeutic strategy against cancer, with documented long-term remissions treating refractory metastatic cancers. However, only a subset of patients (10-40%) have meaningful responses^1,2^. Cancer-cell intrinsic features such as somatic genetic alterations activating specific oncogenic pathways^3,4^, the mutational load^5,6^, the presence and degree of microsatellite instability^7^, and aneuploidy^8^ can differentially influence the responsiveness of the immune system to cancer^9–11^. Modifiable host factors such as the microbiome may also modulate the immune system and affect response to immunotherapy^12,13^.

Tumor intrinsic characteristics influence tumor-immune microenvironment (TIME) composition ^11,14,15^, which itself determines ICI responsiveness^16^. Despite the universe of possible tumor intrinsic characteristics encompassed by all of human malignancy, human tumors can be categorized into a limited number of immune “subtypes” or “archetypes” with prognostic clinical utility^11,17^. While tumor intrinsic features and tumor immune subtypes are important, neither have been proven to determine ICI response. Thus, tumor extrinsic factors such as the tumor host’s germline genetics must play a significant role in ICI-induced anti-tumor immune responses. Supporting this notion, germline genetic variation has a significant impact on virtually every clinically relevant immunologic trait, including autoimmunity, transplant rejection, response to microbial challenges, and cancer susceptibility^18–22^. Since ICI therapeutics exert their effects through an individual’s immune system, it is only rational to assume that genetic variation in host genes participating in immune responses would alter the TIME, and ultimately, modulate anti-tumor responses to ICI treatment.

Prior work suggests that germline genetic variation associated with individual immune genes can alter immunotherapeutic responses. Candidate gene studies of patients on immunotherapy have reported association between various aspects of responsiveness to treatment (CTLA-4 or PD-1 blockade) and polymorphisms in immune-related genes such as IRF5^23^, CCR5^24,25^, CTLA4^26^, HLA^27^, Fc-gamma receptor (FcgR)^28^, IL2, and IL21^20^. The effects of these polymorphisms can be significant. For example, Chowell, *et al.*^27^ found that ICI-treated patients carrying the HLA-B62 supertype had poor survival whereas the HLA-B44 supertype conferred extended survival despite not influencing survival in non-ICI-treated individuals. Naranbhai, *et al.*^29^ extended this finding by showing HLA-A*03 was associated in an additive manner with reduced overall survival after ICI treatment in over 3,000 patients with several types of advanced cancers.

Using a panel of 25 SNPs associated with autoimmune diseases in genome-wide association studies, Chat, *et al.*^20^ examined 436 melanoma patients undergoing ICI treatment. They observed that rs17388568, a risk variant for allergy, colitis, and type 1 diabetes that is located near genes encoding the cytokines IL2 and IL21, was associated with increased anti-PD-1 (aPD1) responses. These studies strongly suggest that germline genetic variations have an impact on immunotherapeutic responses. Unfortunately, these types of studies are often plagued by small sample sizes, involve heterogenous treatments, diseases and patient groups, and examine limited genomic markers (i.e., using a candidate gene/SNP approach), which are experimental conditions prone to generating false positive leads and have limited capacity for discovery of ICI response-modulating genes not already associated with immune traits.

More recently, several groups have leveraged whole genome data to assess the effect of genetic variation on immunological phenotypes with an impact on cancer outcomes. Khan, *et al*.^30^ found that a high polygenic risk score (PRS) for autoimmune conditions (psoriasis and vitiligo) was associated with improved overall survival in patients with bladder cancer treated with atezolizumab (anti-PD-L1). Similarly, the same investigators showed that a high PRS for autoimmune hypothyroidism was associated with lower risk of death in triple-negative breast cancer patients treated with immunotherapies^31^. Other studies have taken more granular approaches to understanding the influence of host genetics on the specific configuration of the TIME. For example, Sayaman, *et al*.^32^ performed a mixed model-based heritability analysis of 139 TIME traits, finding that approximately 25% were heritable, and in some cases, they were able to pinpoint candidate sets of genetic variants associated with traits. Importantly, more refined analyses showed that trait heritability was partially dependent on which of six previously defined tumor immune subtypes^11^ each tumor matched. Pagadala, *et al.*^33^ took a related approach focusing on germline variants having dual associations with TIME characteristics (primarily gene expression levels) and cancer outcomes. This study went a step further by using TIME gene expression quantitative trait loci (QTL) genes to construct PRS modeling cancer risk, overall survival, and immunotherapy response, showing that the latter PRS could discriminate responders from non-responders in independent cohorts.

Nevertheless, even with larger patient cohorts, the major challenge to the mapping of host genes involved in ICI responsiveness remains the genomic heterogeneity inherent to each patient’s unique tumor, even amongst patients with the same type of cancer (i.e. colon cancer patients). Each tumor is derived from an individual’s unique genome yet has acquired further genetic change during neoplastic transformation and immunoediting, therefore the interactions between tumor and host immune system are unique to each patient. These complexities preclude precise identification of ICI response-modulating genes using standard genome wide association (GWAS) approaches in clinical situations.

To address these problems, we have leveraged the inbred nature of the genetically variable Collaborative Cross (CC) mouse genetic resource and an F1 breeding strategy to implant tumors having a fixed genomic configuration (the same tumor model) into mice with variable – yet reproducible – host genetic backgrounds (different CC strains). In this study, we demonstrate that host genetics significantly modulates ICI responses across several distinct tumor models. Moreover, we report four distinct QTL that both additively and epistatically influence response to aPD1 immunotherapy in the context of a colon carcinoma model. Using a systematic approach, we prioritized candidate genes within these QTL and we demonstrate that a transcriptional biomarker outperforms other metrics commonly used in predicting overall survival amongst cohorts of cancer patients treated with aPD1 therapeutics. Furthermore, this transcriptional biomarker showed uniquely high prognostic value for survival in ICI-treated patients whose tumor immune subtypes matched the subtype of our mouse system. Together, these results demonstrate the power of our system for unbiased discovery of genes influencing ICI response and suggest that the quest for the identity of such genes will be aided by well controlled, immune subtype-matched murine models recapitulating key aspects of the TIME in specific and distinct immune subtypes of human cancers.

## Methods

### Contact for Reagents and Resource Sharing

Further information and requests for resources and reagents should be directed to and will be fulfilled by the Lead Contact, Edison Liu.

### Cell culture and DNA isolation

Among the four murine cancer cell lines, CT26 and EMT6 were obtained from ATCC (https://www.atcc.org), AT3 was purchased from Sigma (https://www.sigmaaldrich.com) and MC38 was a gift from Dr. Marcus Bosenberg at Yale School of Medicine, CT USA. AT3 cells were cultured in DMEM--High Glucose medium containing L-glutamine and sodium pyruvate (Sigma Cat. No. D6429) + 1X non-essential amino acids + 15 mM HEPES (7.5 ml of 1 M HEPES for 500 ml media) + 1X β-mercaptoethanol (Cat. No. ES-007-E) + 10% FBS and 1% P/S. CT26 cells were cultured in RPMI1640 (+ 2 mM L-Glutamine) + 10% FBS and 1% P/S. EMT6 cells were cultured in Waymouth (+ 2mM L-glutamine, Gibco #11220035) + 15% FBS and 1% P/S. MC38 cells were cultured in DMEM/F12 (Gibco #11330032) + 1% non-essential amino acids (Gibco #11140050) + 10% FBS + 1% P/S. High-molecular weight DNA and RNA was extracted from frozen cell pellets using AllPrep DNA/RNA/miRNA Universal kit (Qiagen) according to the manufacturer’s protocol.

### Whole genome sequencing

Genomic DNA libraries of 450bp insert size were prepared using the TruSeq DNA PCR-free Library Preparation Kit (Illumina) according to manufacturer guidelines and sequenced to at least 30x coverage by the New York Genome Center (NYGC). The sequencing data for all four cell lines was processed and aligned to the mouse genome (mm10) through NYGC’s variant somatic pipeline v6 (https://bioinformatics.nygenome.org/wp-content/uploads/2019/06/SomaticPipeline_v6.0_Human_WGS.pdf). In brief, reads were aligned to the mouse genome (mm10) using the Burrows-Wheeler Aligner (BWA)^34^. Structural variant (SV) calls were generated using three different tools (Crest^35^, Delly^36^, and BreakDancer^37^), and high confidence events were selected when called by at least two tools and by requiring split-read support. Single Nucleotide Variants were called by muTect^38^, Strelka^39^ and LoFreq^40^. We considered high confidence mutations as those identified by all three tools. The reference C57BL/6 genome was used as a matched normal for MC38 and AT3 while the BALB/cJ genome was used for CT26 and EMT6. Somatic mutations were considered those unique to each cell line and absent from an in-house generated panel of normal samples (PON) comprising 11 normal mouse genomes. In addition, high-confidence SVs were identified by filtering out SVs detected in an in-house cohort of 63 mouse cancer genomes and those only detected by one SV caller or not supported by split reads, to reduce possible germline events and sequencing artifacts. Tandem duplicator Phenotype (TDP) was assessed as described elsewhere^41^.

### Mutational signature analysis and tumor mutational burden (TMB)

We ran deconstructSigs (v1.8.0)^42^ on the high confidence somatic SNV call set within autosomes to estimate the contribution of known COSMIC mutational signatures (v1-2013)^43^ in the tumor sample. TMB was defined as the number of somatic, coding, SNVs and short indels per megabase of genome examined^44^. Mutations that are present in dbSNP (dbSNP 142)^45^ were excluded from the analysis.

### Tumor growth and ICI efficacy studies

To measure tumor growth and assess ICI efficacy, tumor cell lines were injected into female mice aged 8-12 weeks. For MC38 and CT26, 0.5 million cells were injected subcutaneously seven days before ICI dosing. For AT3 and EMT6, cells were injected orthotopically into mammary fat pads at a dosage of 0.3 million (EMT6) or 1 million (AT3) cells either seven (EMT6) or ten (AT3) days before ICI dosing. During the dosing period, either isotype control (Bioxcell, 2A3) or aPD1 antibody (Bioxcell, 29F.1A12) was injected intraperitoneally at 200ug per mouse every three days culminating in a total of four doses. Throughout the study, body weights, tumor measurements, and clinical observations were performed three times per week. Mice were removed from a study and euthanized if animal care staff noted tumor ulceration, body weight loss >20%, or any other veterinary care issue. For single dose studies, measurements of body weight and tumor size were performed daily after tumor cell injection. When tumors reached a size of 75-110mm^3^, a single dose of isotype control or aPD1 antibody was administered and the mice were euthanized 48 hours later.

### Quantification of tumor growth rates

In order to calculate a quantitative measure of the mean and variance in response to aPD1 immunotherapy, we devised a per-mouse metric to measure tumor growth and regression after treatment. We developed this metric within the framework of the treatment/control tumor volume ratio, which is widely used in tumor xenograft studies. Hather, *et al*.^46^ showed that a summary metric they called the rate-based T/C is a straightforward and powerful method for estimating treatment efficacy in such a study. This method makes the assumption that tumor volume follows an exponential growth trajectory and fits a linear regression model to derive a growth rate for each tumor from the logarithm (base 10) of tumor volumes. The rate-based T/C is computed as Rate-based *T* / *C* = 10^(*μ*_T_−*μ*_C_)×21 days^, where *μ*_T_ is the mean of the tumor growth rates for the treatment group, and *μ*_C_ is the mean of the tumor growth rates for the control group^46^. In effect, this method compares the slopes of log-transformed tumor volume measurements through the study period and computes the ratio of these numbers as a measure of growth inhibition in the treated group compared to the control group. We adapted the rate-based T/C to apply to our syngeneic host model, computing these quantities separately within each genetically distinct line (e.g. CCF1, B6, BALB). Rather than computing group means for the treated and control groups, we computed per-mouse T/C ratios by extracting tumor growth curve slopes for each individual mouse and normalizing by dividing by the mean slope in the same line’s control group. Therefore, our metric is computed using the formula: per-mouse rate-based *T* / *C* = 10^(*b_i_*−*μ*_C_)×21^, where *b_i_* denotes the tumor growth rate of mouse i. When computing slopes from log-transformed tumor volumes, we considered data starting from the first day of ICI dosing (e.g. day 7 for MC38), and omitted measurements of tumors below detectable size (because the logarithm of zero is undefined). For tumors that initially grew but quickly regressed to zero after a single non-zero measurement, we inserted a small value for tumor growth slope in order to capture this responsiveness to ICI. Specifically, we used the 10^th^ quantile of all (log-transformed) tumor growth slopes, computed separately in each treatment group (isotype or aPD1). For QTL mapping, we mapped the mean of the logarithm of per-mouse rate-based T/C computed for each CCF1 line.

Finally, we found that a small number of lines appeared to have an enrichment in the aPD1 treated group of mice carrying tumors that never grew (complete responder*; CR* tumors). Because this enrichment was specific to the aPD1 treatment group, it likely represented tumors that were super responders but could not be registered using the RTC methology. We used the binomial exact test to quantify how unusual the enrichment of CR* tumors was in each CCF1 line. Two CCF1 lines (CC75 and CC68) showed significant enrichment of CR* tumors in aPD1 compared to isotype control group. In order to include this component of aPD1 response in our mapping, we used the large sample binomial approximation to obtain an approximate Z-score for this test and computed a weighted average with the RTC metric (first converting to a Z-score) using weights 20% and 80% because approximately 20% of lines had >1 CR* in the aPD1 treatment group. The results of mapping this combined trait were substantially similar to mapping of RTC alone, and the peak on Chr15 (main text) had a marginally more significant LOD score (data not shown).

For CCF1N1 mapping studies, we could not compute the per-mouse rate-based T/C described above because each mouse was genetically unique and there was no isotype control group for normalization. Instead, we mapped the un-normalized quantity (slope of the log-transformed tumor growth curve). We also mapped two additional traits consisting of discrete classifications of response. First, we mapped the binary indicator of whether (or not) each mouse’s tumor was a complete responder (CR) to treatment. Second, we mapped a more granular classification of tumor response, namely whether each tumor was a non-responder, partial responder, CR, or CR*. In the main text we present results of mapping the binary CR indicator because this method was simple and utilized all mice. Nevertheless, all QTL we report in the main text were prominent and statistically significant in at least two of the three traits mapped.

### Quantitative trait locus mapping

To map quantitative trait loci (QTL) that harbor genetic elements mediating variation in the response to aPD1 immunotherapy, we considered our CCF1 lines as a genetic mapping population. We genotyped one female and one male mouse of each CC strain using the GigaMUGA array^47^. We compared our data to previously reported strain genotypes derived from mice sourced from a different institution^48^ and found high agreement, with the closest match to each strain having the same strain designation. We used in-house female genotypes for QTL mapping since we performed aPD1 testing in female mice. To conduct QTL mapping we utilized the R/qtl2 package^49^ using the “genail” cross type. To initialize mapping we followed https://github.com/rqtl/qtl2data/blob/master/CC/R/convert_cc_data.R and obtained the necessary funnel codes describing the breeding scheme for this genetic resource population from Shorter, *et al.*^50^. We mapped the specific phenotypes described above (“Quantification of tumor growth rates”) and used a permutation test to establish statistical significance as implemented in the qtl2 function scan1perm, with the following parameters: 1000 permutations and significance threshold 0.05.

### Prioritization of candidate genes within QTL

To prioritize candidate genes with QTL, we developed an algorithm to define a modest set of genes that strongly track with ICI response in the MC38 system and in human cohorts. First, we defined ICI response-associated regions within each mouse mapping population and set confidence boundaries on the location of the QTL using 95% Bayesian credible intervals^51^. We used MC38 tumor bulk RNA-Seq samples to define genes that are expressed in tumor samples (defining “not expressed” as <5 reads across all samples). To identify genes that might be associated with response in human tumor transcriptomes, we leveraged TCGA data available through the Xena browser^52^. Since the MC38 cell line is derived from a colon carcinoma and we studied response to aPD1 immunotherapy, we subset TCGA samples to those classified as colon adenocarcinoma, lung adenocarcinoma, lung squamous cell carcinoma, and skin cutaneous melanoma (lung and skin cancers being commonly treated with aPD1 therapeutics). Because no measure of immunotherapy response is available for TCGA cohorts, to allow us to leverage these relatively large transcriptome datasets we computed the ratio CXCL9/SPP1, which has been associated with favorable tumor microenvironment^53^. We computed the Spearman correlation of human orthologs of each mouse gene to the CXCL9/SPP1 ratio in the four TCGA cohorts chosen. Next, we classified each gene according to whether it resides within the QTL credible interval and whether the gene was differentially expressed in MC38 bulk tumor RNA-Seq samples (responder vs. non-responder samples). To prioritize top candidate quantitative trait genes located within QTL, we used genes outside the QTL to construct an empirical null distribution of the absolute value of the correlation to CXCL9/SPP1 ratio. For each TCGA cohort we separately defined a threshold as the top quartile value for this correlation (from the empirical null distribution) and extracted genes exceeding this correlation that were within QTL and differentially expressed in murine tumor RNA-Seq. Subsequently we ranked genes according to the strength of the absolute correlation to CXCL9/SPP1. We called this approach the Cross-Species TIME algorithm.

### Testing prognostic value of transcriptional biomarkers in immunotherapy-treated cohorts

To determine whether prioritized quantitative trait genes can be used to form a transcriptional biomarker with prognostic value for immunotherapy response, we turned to cohorts of immunotherapy-treated patients with both transcriptional profiles and overall survival recorded. We obtained data from immunotherapy-treated patient samples of seven cancer types^54,55^. After filtering to patients treated with aPD1 therapeutics and having available pre-treatment transcriptional profiles, we assembled a dataset of 551 patients from seven studies.

To compute prognostic score for immunotherapy response, we first split the data by study and Z-transformed gene expression values. To calculate a score for each individual in the study, we multiplied each gene by a weight (−1 or +1 according to whether the gene was negatively or positively correlated with CXCL9/SPP1 transcript ratio in TCGA data) and summed all weighted Z-transformed gene expression values for the individual. To normalize across samples that could have different amounts of missing data, we multiplied this score by the reciprocal of the fraction of genes that were missing data in each individual. We used the survival and survminer R packages to conduct all survival analyses. When plotting survival curves we assigned individuals into “score high” or “score low” groups according to whether that individual’s score was above or below, respectively, the median score of all individuals in the study. To test the predictive value of our transcriptional biomarker on survival, we used a stratified model where overall survival was modeled as a function of the normalized biomarker score and stratified by study. We used a likelihood ratio test to determine the significance of the inclusion of the normalized biomarker score in the model.

### Digestion of Tumor Samples

Mice were euthanized via cervical dislocation, tumors were excised and separated from surrounding skin using a razor blade. Tumors were split into 2mm wide strips (if necessary) and were submerged in 10% DMSO (Millipore Sigma) in DMEM (ThermoFisher Scientific) and gradually frozen in isopropanol at −80°C overnight before storage in liquid nitrogen. For analysis, samples were thawed to 37°C in a water bath and rinsed in “FACS buffer” (1x Ca^2+^ and Mg^2+^ deficient HBSS (ThermoFisher Scientific) supplemented with 1% BSA (Research Products International)). The samples were thoroughly minced into <0.5mm pieces using a sterile razor blade and digested in 6mL “digest solution” (1x HBSS with Ca^2+^ and Mg^2+^ HBSS (ThermoFisher Scientific) supplemented with 1% BSA, Collagenase VIII (Millipore Sigma) at 2mg/mL, and DNase I (Millipore Sigma) at 50ug/mL). Samples were incubated at 37°C and shaken at 250RPM for 30 minutes. Samples were Q.S. to 20mL with FACS buffer, filtered through a 100µm strainer, and centrifuged at 500g at 4°C for 10 minutes. After decanting digest solution, samples were resuspended in 5mL Hybri-Max Red Blood Cell Lysis (Millipore Sigma) for 7 minutes before Q.S. to 30mL with FACS buffer and centrifuged again before sample processing.

### Cell Staining for FACS

Cells were stained for 30 minutes on ice in the dark with an MHC Class I Tetramer (PE) (KSPWFTTL, NIH Tetramer core) recognizing an MC38 antigen-specific T-cell receptor (TCR), PE-CF594-CD3e (clone 145-2C11; BD Biosciences, San Jose, CA), PerCP-Cy5.5-CTLA4 (UC10-4B9; Biolegend, Inc., San Diego, CA), PE-Cy7-F4/80 (BM8.1; Biolegend, Inc., San Diego, CA), APC-MHCII (M5/114.15.2; Biolegend, Inc., San Diego, CA), AF700-CD206 (C068C2; Biolegend, Inc., San Diego, CA), AFC-AF750-Ly6C (HK1.4; Biolegend, Inc., San Diego, CA), BV421-PD1 (RMP1-30; Biolegend, Inc., San Diego, CA), BV510-CD4 (GK1.5; Biolegend, Inc., San Diego, CA), BV570-CD45 (30-F11; Biolegend, Inc., San Diego, CA), BV605-PDL1 (MIH6; Biolegend, Inc., San Diego, CA), BV650-CD11b (M1/70; BD Biosciences, San Jose, CA), BV711-CD8a (53-6.72; BD Biosciences, San Jose, CA), BV786-Ly6G (1A8; Biolegend, Inc., San Diego, CA), FC Block (anti-CD16/32 (clone 2.4G2; Lienco Technologies, Inc., Fenton, MO)) and Brilliant Stain Buffer Plus (BD Biosciences) in FACS buffer. Samples were washed twice by Q.S. to 1mL and resuspension, centrifuging at 500g for 10 minutes at 4°C, and decanting the liquid. Samples were then resuspended in 250uL 1:1000 YO-PRO (ThermoFisher) in FACS buffer and filtered through a 35µm strainer into FACS tubes on ice in the dark (centrifuge at 500g for 20 seconds, if necessary, to pass cells through strainer). DAPI (ThermoFisher Scientific) was added and each sample was vortexed for three seconds before immediate detection on a FACSymphony A5 SE (BD Biosciences). Samples were analyzed with FlowJo software (BD Biosciences).

### Bulk RNA-Seq data generation and analysis

For bulk RNA-Seq samples, we harvested tumors and froze in DMEM/10% FCS with 10% DMSO. We harvested RNA using the RNeasy (QIAGEN) RNA extraction kit. Poly(A) RNA-seq libraries were constructed using the KAPA mRNA HyperPrep Kit (KAPA Biosystems). Libraries were sequenced 100 bp paired-end on the NovaSeq 6000 (Illumina) using the S4 Reagent Kit (Illumina). We constructed transcriptomes for each CC founder strain using GENCODE vM23 annotations by using g2gtools v0.2.9 (https://github.com/churchill-lab/g2gtools) to alter sites that differed from the mouse reference genome according to Mouse Genomes Project SNP and indel release version 5^56^. We mapped reads to these transcriptomes using bowtie v1.2.3^57^, followed by converting to EMASE format and using gbrs v0.1.6^58^, which can account for reads mapping to multiple or divergent haplotypes, to quantify multiway expression across the eight founder haplotypes. We used only total expected read counts across each gene for our analyses.

For differential expression testing, we used the DESeq2 package v1.44.0^59^ with default options and following author recommendations. For the heatmap produced in Figure 5D, we performed differential expression testing on all pairwise combinations of groups in our blocking antibody experiment, identified enriched Gene Ontology biological process pathways using log2FoldChange values and the fgsea package v1.30.0^60^, pruned to semantically distinct terms (Wang measure, threshold 0.6) using GOSemSim v2.30.0^61^, and filtered to immune-related pathways by considering only terms descended from the following list of immune-related GO terms: “immune response”, “immune system process”, “response to cytokine”, “regulation of immune system process”. For the principal components analysis (PCA) depicted in Figure 5E, we started with bulk RNA-Seq samples from (1) genetic responders and non-responders (2) our blocking antibody experiment, normalized gene expression using edgeR v4.2.0^62^, leveraged the ComBat method^63^ to minimize batch effects between these two sets of samples, and using the batch-adjusted matrix for PCA. For deconvolution of bulk RNA-Seq samples in the blocking antibody experiment, we used mMCP-Counter^64^ with default parameters.

### Tumor immune subtype classification

In order to classify the immune subtypes of tumors, we used the ImmuneSubtypeClassifier R package (https://github.com/CRI-iAtlas/ImmuneSubtypeClassifier)^65^, which predicts subtype membership based on the classifications proposed by Thorsson et al.^11^. For human samples we applied this algorithm to normalized gene expression of each sample according to package documentation. For mouse samples we converted gene symbols to their human counterparts using babelgene^66^, summing up counts for cases where a single human gene mapped to multiple paralogs in the mouse. We then applied ImmuneSubtypeClassifier as for human samples.

### Single cell RNA-Seq data generation

Dissociated tumor cells from responder and non-responder mice strains were incubated with FcR blocking solution, then stained with cell hashing antibodies (TotalSeq-C anti-human Hashtag (HTO) C0301-C0304, BioLegend) and with BV570-CD45 (clone 30-F11), Brilliant Stain Buffer Plus in FACS buffer in the dark on ice for 30 minutes. After a FACS buffer rinse, samples were then resuspended in 1:1000 YO-PRO in FACS Buffer and filtered through a 35µm strainer into FACS tubes on ice in the dark (centrifuge at 500g for 20 seconds, when necessary, to pass cells through strainer). DAPI was added and each sample was vortexed for three seconds before immediate sorting (FACSAria, BD Biosciences). Combined samples stained with different hashtags were sorted into the same collection tube for single cell RNA-Seq. From the total viable cells in each sample (DAPI^−^YOPRO1^−^), 90% CD45^+^ and 10% CD45^−^ cells were sorted into 5μL FBS per collection tube, keeping the concentration from each sample equal in each tube if possible. Cell viability was assessed with Trypan Blue on an automated cell counter (Countess II, ThermoFisher), and up to 20,000 cells (∼5,000 cells from each hashtagged sample) were loaded onto one lane of a 10x Genomics chip, then cells and gel beads were portioned in a Chromium X instrument (10x Genomics). Library preparation was performed using NextGEM single cell 5’ version 2 chemistry and according to the manufacturer’s protocol (10x Genomics, CG000330). cDNA and libraries fragments were profiled by automated electrophoresis (Tapestation 4200, Agilent) and High Sensitivity D5000 reagents and fluorometry using High Sensitivity 1x dsDNA reagents (Qubit, Invitrogen) and the presence of Illumina adapters was verified via qPCR (QuantStudio 7 Flex, Applied Biosystems). Libraries were then pooled using a ratio of 90% gene expression library and 10% HTO library before sequencing; each gene expression-HTO library pair was sequenced at 15% of an Illumina NovaSeq 6000 S4 v1.5 200 cycle flow cell lane, with a 28-10-10-90 asymmetric read configuration, targeting 10,000 barcoded cells with an average sequencing depth of 75,000 reads per cell.

### Fresh frozen Visium spatial transcriptomics

After resection, each tissue was blotted dry with an RNase-free laboratory towel then immediately submerged in fresh isopentane chilled over liquid nitrogen for approximately one minute and transferred to a cryovial pre-cooled on dry ice for long term storage at −80C. Tissue was subsequently submerged in Optimal Cutting Temperature compound (OCT) in a prechilled cryomold using prechilled forceps, and the cryomold was immediately placed on powdered dry ice and allowed to completely freeze. OCT blocks were stored at −80C until quality control and processing.

For spatial transcriptomics, five 10µm sections from each block were used for total RNA extraction (RNeasy Mini Kit, Qiagen) and determination of RNA integrity (RINe score) using an automated electrophoresis (Tapestation 4200, Agilent) and High Sensitivity RNA ScreenTape. Blocks with RINe greater than 7 were optimized for ideal permeabilization time following the vendor protocol (10x Genomics, CG000238). Sections from four tissue blocks were placed on Visium Gene Expression slide for H&E staining and brightfield imaging via NanoZoomer SQ (Hamamatsu), block-specific tissue permeabilization, mRNA capture, and subsequent library generation per the manufacturer’s protocol (10x Genomics, CG000239). Libraries were quantified by automated electrophoresis (Tapestation 4200, Agilent) and High Sensitivity D5000 reagents and fluorometry (Qubit, Invitrogen) and the presence of Illumina adapters was verified via qPCR (QuantStudio 7 Flex, Applied Biosystems). Libraries passing quality control were pooled for sequencing on an Illumina NovaSeq 6000 200 cycle S4 flow cell using a 28-10-10-90 read configuration, targeting 100,000 read pairs per spot covered by tissue. Illumina base call files for all libraries were converted to FASTQs using bcl2fastq v2.20.0.422 (Illumina). Whole Visium slide images were uploaded to a local OMERO server. For each capture area of the Visium slide, a rectangular region of interest (ROI) containing just the capture area was drawn on the whole slide image via OMERO.web and OMETIFF images of each ROI were programmatically generated using the OMERO Python API.

### Single cell RNA-Seq analysis

Illumina base call files for all libraries were converted to FASTQs using bcl2fastq v2.20.0.422 (Illumina) and FASTQ files associated with the gene expression libraries were aligned to the GRCm38 reference assembly with vM23 annotations from GENCODE (10x Genomics mm10 reference 2020-A) using the version 6.1.1 Cell Ranger count pipeline (10x Genomics). We carried out analyses of processed scRNA-Seq data in R version 4.3.3^67^, using Seqgeq software (BD Biosciences), and the R package Seurat version 5.0.1^68^. We quantified expression across 32,285 genes, of which 26,938 were expressed in at least one cell. After filtering out cells with >15% mitochondrial gene expression or >45% ribosomal gene expression, we obtained data from 93,126 cells passing quality control steps. We normalized using log normalization and selected 2,000 variable features. We scored cells according to cell cycle following documentation available with Seurat. We scaled/centered feature expression after regressing out the total read count, percent mitochondrial gene expression, percent ribosomal gene expression, and cell cycle phase of each cell. We reduced dimensionality using principal components analysis (PCA) and used harmony v0.1^69^ with 37 PCs using option theta=1 to correct for batch processing date of our scRNA-Seq libraries. for cell clustering. We used the shared nearest neighbor-based modularity optimization algorithm implemented in Seurat to cluster cells, using resolution 0.1, and projected into UMAP space using 37 PCs. For subclustering of monocyte/macrophages, we reanalyzed these cells and clustered with 26 PCs and using clustering resolution 0.6. For T/NK cells we performed a similar analysis but using 19 PCs and clustering resolution 0.3.

### Spatial transcriptomics analysis

FASTQ files and associated OMETIFF corresponding to each capture area were aligned to the GRCm38 reference assembly with vM23 annotations from GENCODE (10x Genomics mm10 reference 2020-A) using the version 1.3.1 Space Ranger count pipeline (10x Genomics). We carried out analyses of processed spatial transcriptomics data in R version 4.3.3^67^, using Seqgeq software (BD Biosciences), and the R package Seurat version 5.0.1^68^. We normalized spot gene expression using the SCTransform algorithm implemented in Seurat. To test for colocalization of cell types: cytotoxic T lymphocytes (CTLs) with dendritic cells (DCs) or *Cxcl9^+^* macrophages) we computed binary indicators of nonzero expression of the marker genes mentioned (main text) for each spot and deemed two cell types as colocalized within a spot if markers of both cell types were expressed in that spot.

## Results

### QUANTIFYING HOST HERITABILITY (H^2^) OF ICI RESPONSE

To establish an experimental system allowing for the identification of host genetic factors influencing ICI response, we took advantage of common tumor engraftment models and the Collaborative Cross (CC) mouse genetic reference population^59^. Our initial goal was to calculate the heritability of the response to aPD1 therapy across these distinct tumor lines to ascertain whether host genetics impacts ICI responsiveness. The major challenge for tumor transplantation models in genetically diverse backgrounds is allogeneic tumor rejection. To overcome this impediment, we used an F1 hybrid approach whereby genetically diverse inbred CC strains were crossed to the inbred genetic background of the transplanted tumor cell line (**Figure 1A**)^70^. We chose multiple murine tumor cell lines from two genetic backgrounds, C57BL/6J (B6) and BALB/cJ (BALB), covering both mammary gland and colon cancers: MC38 (colon; B6), CT26 (colon; BALB), AT3 (mammary gland; B6), and EMT6 (mammary gland; BALB). These cell lines were selected because they have been shown to be responsive to ICI treatment^71–73^ and because they provided a balanced mix between different tissues of origin (mammary and colon) and genetic backgrounds (BALB and B6). Our initial goal was to calculate the heritability of the response to aPD1 therapy across these distinct tumor lines to ascertain whether host genetics broadly impacts ICI responsiveness.

**Figure 1:**
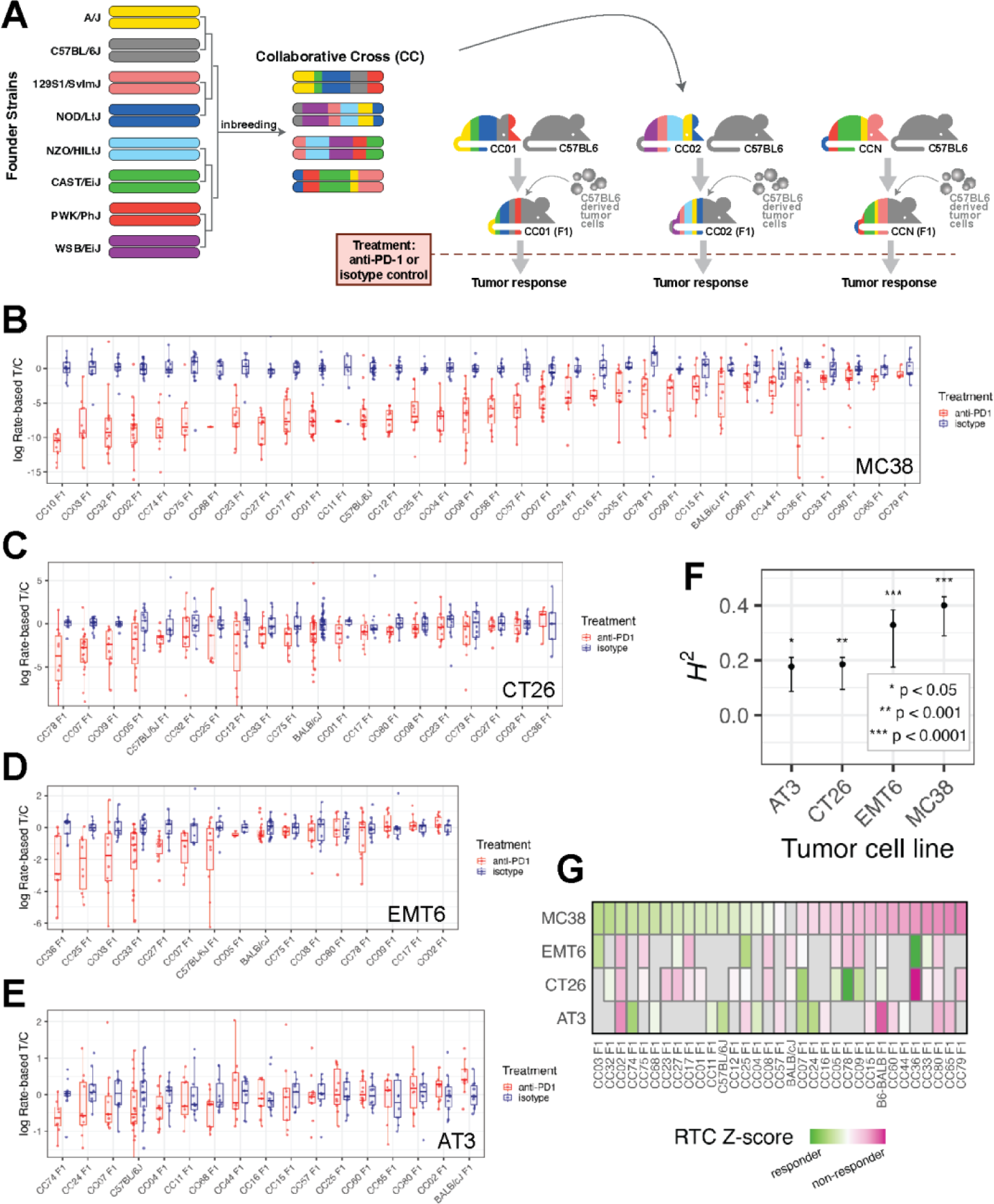
A system for quantifying ICI response across variable host genetics and its application to four implantable tumor models. (A) Schematic overview of our mouse CC-based system for measuring ICI response in diverse but replicable genetic backgrounds using a tumor with fixed genetic background. (B-E) Boxplots showing our measure of ICI response, the per-mouse rate-based tumor/control (RTC) computed across strains implanted with each tumor cell line model. Blue points and boxes display RTC for isotype control mice which are approximately centered at zero for all strains, while red points/boxes display values for aPD1-treated mice. A large difference between red and blue boxes for a particular strain would indicate a strong response to ICI. (F) Significant broad-sense heritability (*H*^2^) of ICI response was seen for each implantable tumor cell line model, computed using data from all CCF1 lines profiled with that model. (G) Heatmap of median log(RTC) values standardized across each tumor model and plotted across all strains profiled in at least two tumor models show differential impact of the CCF1 crosses on aPD1 response depending on the tumor model used.

First, we constructed host recipients for the MC38 colon cancer line. These recipients were F1 offspring of crosses between B6 and each of 32 inbred CC lines (F1 offspring referred to here as “CCF1”, **Figure 1A and Table S1**). The CC is a multiparent, recombinant inbred strain panel derived from eight inbred founder strains segregating ∼40 million genetic variants^74–76^. Each CCF1 tumor recipient mouse carried maternal ancestry from a CC strain together with paternal ancestry from B6. The advantage of the CC resource is that each line carries a fixed set of genetic variants present on haplotype blocks segregating at a frequency of approximately 1/8 (eight founder strains with evenly distributed genetic contributions). This approach produced a set of CCF1 strains with a high degree of well-defined genetic diversity matched with the experimental replicability of inbred strains. We created a panel of CCF1 recipients for the AT3 cell line in a similar manner, as well as crossing CC strains to BALB to create CCF1 recipients for the CT26 and EMT6 (i.e. BALB-derived) lines.

### VARIATION IN ICI RESPONSE IS HERITABLE IN GENETICALLY DIVERSE MICE

To measure ICI response in individual strains of mice, we developed a method to quantify the magnitude with which ICI treatment retards tumor growth in comparison to treatment with isotype control antibody in genetically identical mice called the per-mouse rate-based tumor/control^46^ (RTC; see Methods). The RTC metric (measure of ICI response) assumes tumors grow exponentially and utilizes linear regression modeling of the logarithm of tumor size to derive an estimate of tumor growth rate that is robust to statistical noise and missing data. We found that the variation in RTC within a CCF1 strain is less than that observed between strains (e.g. **Figure 1B-E**).

When we applied this approach to the MC38 system, we observed a broad range of aPD1 responses spanning from individual CCF1 strains that exceeded the parental (B6) response with nearly all complete responders (CC02 F1), to strains with no discernable response (CC11 F1) (**Figure 1B**). Across 34 lines (including 32 genetically diverse CCF1 lines, inbred B6, and B6/BALB F1 mice), quantitative genetic analysis revealed that ∼40% of the total variation in ICI response in the MC38 model is attributable to genetic background in our CCF1 system (broad-sense heritability *H*^2^ = 0.40; *P* < 1×10^−4^) (**Figure 1F**).

Given this positive result, we then scanned >15 CCF1 crosses using each of the other three murine cancer models (**Figure 1C-E**) and found significant heritability for each (**Figure 1F**): 0.33 for EMT6 (*P* < 1×10^−4^), 0.19 for CT26 (*P* = 2×10^−4^), and 0.18 for AT3 (*P* = 0.017). Interestingly, each tumor model had a distinct set of responder and non-responder CCF1 strains (**Figure 1G**). For example, CC02F1 mice are good responders in the MC38 system, but poor responders for the other tumor models, and while CC78F1 mice show a good response in the CT26 model, they are non-responders in the EMT6 and MC38 models. This suggests that tumor intrinsic factors influence how host genetics mediates responsiveness to ICI.

The microbiome has been reported to alter the efficacy of ICI therapeutics^77^. To assess this possibility, we tested two aPD1 responder CCF1 lines (CC02 and CC75) for response with and without pre-treatment with a broad-spectrum cocktail of antibiotics using a protocol that substantially reduces the gut microbiome and its effects on other phenotypes^78^. Importantly, we found no diminution of the ICI response to MC38 tumors (**Figure S1A,B**). These results strongly suggest that ICI responses in our MC38 CCF1 system are a direct effect of host genetics and not influenced by the microbiome.

Though previous reports of the sequence characteristics of CT26, MC38, and EMT6 exist^79,80^, we re-sequenced all four cell lines to compare mutational burden and other relevant genomic characteristics. When the two cell lines with the highest heritability (MC38 and EMT6) were compared to the two with the lowest H^2^, no distinguishing features emerged. MC38, a B6 derived colon carcinoma, had the highest tumor mutational burden (TMB = 112 mutations per Mb; **Figure S1C**) and is MSI (microsatellite instability) positive^81^. EMT6, which had the second highest H^2^, has low TMB (18.5 mutations per Mb) and is a mammary carcinoma line in a BALB genetic background. Surprisingly, given the association of TMB and ICI response, the CT26 cell line had an H^2^ of only 0.19 (**Figure 1C**) despite having the second highest TMB of 92 mutations per Mb (**Figure S1C**). Moreover, no characteristic single base substitution signature or other structural variations distinguished high heritability vs. lower heritability (**Figure S1D**). Thus, genomic analyses of these four tumor models showed no obvious genomic signature linked to aPD1 response.

### IDENTIFICATION OF ANTI-PD-1 RESPONSE QTLS AND THEIR EPISTATIC INTERACTIONS

The MC38 CCF1 model exhibited the highest estimated heritability of ICI response (**Figure 1F**) and the largest dynamic range of responses (**Figure 1B**). We carried out quantitative trait locus mapping using 32 CCF1 strains and identified a locus on mouse Chr15 having significant association with aPD1 response (LOD = 8.9) along with multiple subthreshold peaks (**Figure 2A**). Understanding that the CCF1 approach has limited mapping power due to the structure of this population and because only 32 CCF1 lines were used^82^, we sought to refine these subthreshold peaks using a more powerful mapping design. Therefore, we constructed intercrosses between an ICI responder (CC75) and a non-responder CC line (CC80) and mated the offspring of this CC intercross to B6 female mice (a design which we have termed CCF1N1 mapping, schematized in **Figure 2B**). While paternally-derived CC strain chromosomes are genetically variable, the resulting mice carry maternally-derived chromosomes from an inbred B6 background, which initiates host immunological compatibility with the MC38 model. Moreover, because the paternal chromosomes are derived from a cross between responder and non-responder CC lines, this design guarantees that ICI response-modulating genetic variation segregates in the mapping population.

**Figure 2:**
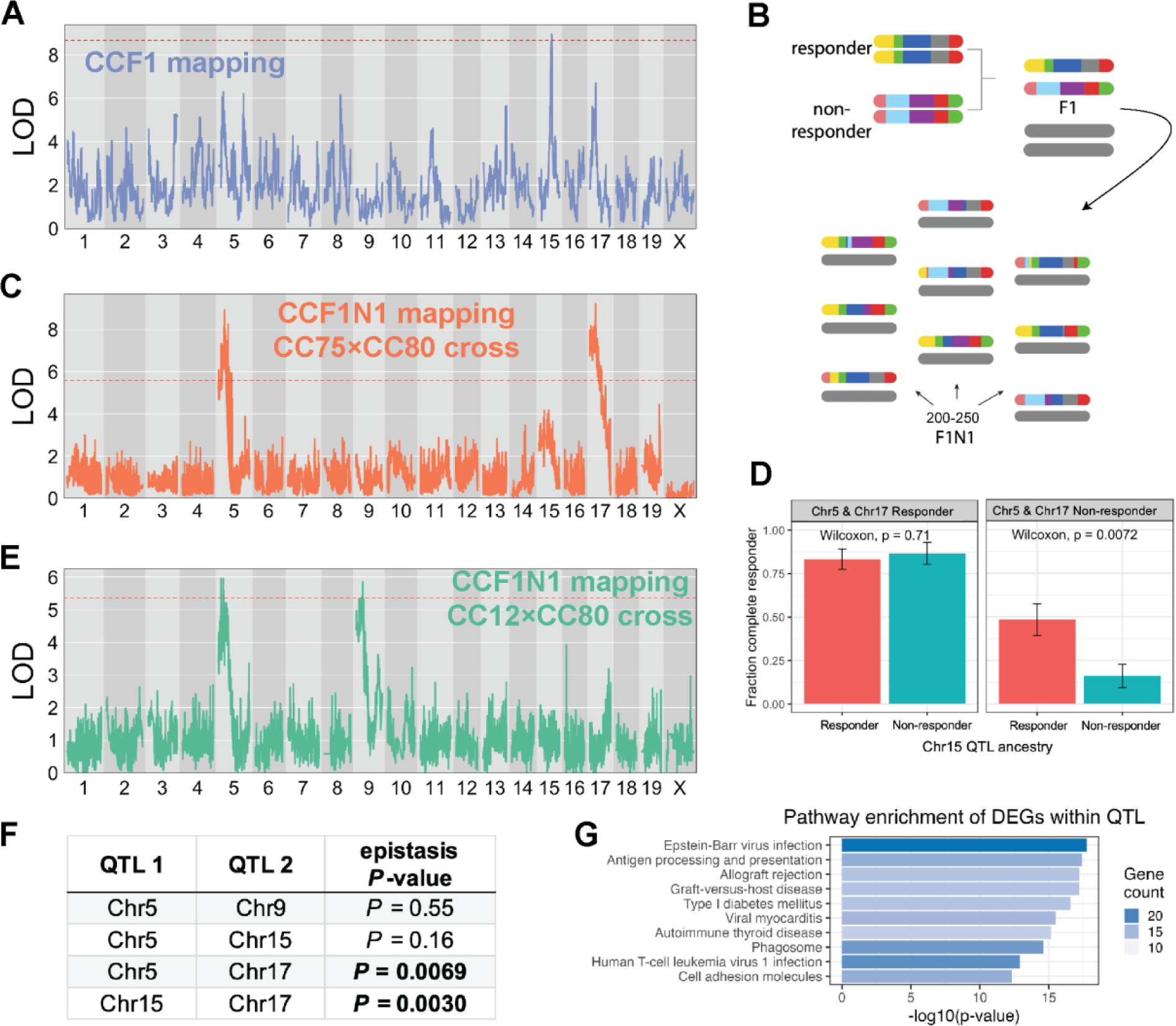
Discovery of ICI response QTL and their epistatic interactions. (A) LOD curve for a genome scan of ICI response in CCF1 lines using the MC38 tumor model. (B) Schematic outlining breeding design to generate a mapping population for CCF1N1 studies. (C) LOD curve for a genome scan of ICI response in CCF1N1 (CC75×CC80) mice using the MC38 tumor model. (D) Illustration of epistatic interactions between QTL in our CCF1N1 (CC75×CC80) mapping population. Three QTL on mChr 5, 15, and 17 were polarized as responder or non-responder using haplotype effect estimates from our CCF1 mapping. The results show that the responder haplotype on Chr15 exhibited an *in vivo* response mainly when both the Chr5 and Chr 17 haplotypes are in the non-responder configuration. (E) LOD curve for a genome scan of ICI response in CCF1N1 (CC12×CC80) mice using the MC38 tumor model. Since the two CC lines are matched for the mChr 17 haplotype, the mChr 17 QTL is absent. (F) Table reporting results of statistical testing of epistatic interactions between QTL (see Methods). Pairs of QTL with significant evidence of interaction are shown with *p-*value in bold. (G) Top 10 most enriched pathways among responder vs. non-responder strain differentially expressed genes that are physically located within one of our four QTL intervals. Bar length is proportional to the −log10(*p*-value) for enrichment, while bar shading is proportional to the number of genes in each annotated pathway contained within our gene set (DEGs within QTL).

In the CCF1N1 design, each mouse is genetically unique and therefore cannot be normalized to an isotype control-treated control group. For QTL mapping in this population (*N =* 249 mice), we mapped either un-normalized RTC or a discrete classification of response (Methods). We identified significant QTLs for aPD1 response on mChr5 and mChr17, which originally appeared in CCF1 mapping as statistical subthreshold peaks (**Figure 2A versus 2C**). While the original CCF1 mapping showed a prominent mChr15 QTL, this peak, while present, lost statistical significance in the subsequent CCF1N1 crosses with the emergence of QTLs on mChr5 and 17. One explanation for this observation is that the magnitude of the mChr15 QTL’s effect depends on the genetic context. We hypothesized that intercross-specific combinations of responder haplotypes in each of these QTLs could significantly modulate the impact of a responder haplotype at a locus under study. Indeed, we observed epistatic interactions between QTLs: the QTL on mChr15 did not affect response when mice carried responder haplotypes at mChr5 and 17 QTL, yet the mChr15 QTL had a strong effect on response in mice carrying non-responder haplotypes at these loci (**Figure 2D**).

This result suggested that additional QTL might be uncovered in specific genetic contexts while being masked by epistatic effects in other contexts (e.g. a different CCF1N1 population). We tested this hypothesis by initiating a CCF1N1 study where we selected lines for the CC intercross that had matched ancestry at the mChr17 QTL. The mChr17 QTL encompasses the MHC locus, which includes MHC class I and II molecules. We asked whether we could identify additional MC38 response-modifying QTLs by generating CCF1N1 mice (*N =* 237) using a responder (CC12) strain and non-responder strain (CC80) with matched ancestry at the mChr17 QTL but segregating variable haplotypes (responder vs. non-responder) at the mChr5 and 15 QTLs. With this approach, as expected, the mChr17 QTL was eliminated while the QTL on mChr5 was detected. However, consistent with our speculation about potential masking of QTL by epistasis, we observed a new and significant QTL on mChr9 (**Figure 2E**).

Because of the structure of the genetic crosses, where multiple QTL segregated in a cross, we could statistically quantify the extent of epistasis between them using logistic regression modeling (**Figure 2F**). For the first CC80×CC75 CCF1N1 experiment, a model including all possible QTL interactions was strongly favored over the null hypothesis of no epistasis (*P* < 0.0001). We did not observe evidence for 3-QTL interaction (*P* = 0.34), pointing to the importance of pairwise interactions between QTL. The mChr15–17 and mChr5–17 QTL interactions were statistically significant while the Chr5–15 QTL interaction was not (**Figure 2F**). An independent examination of the second CC12×CC80 CCF1N1 experiment revealed no epistasis between the Chr5 and Chr9 QTL (**Figure 2F**). Overall, these results demonstrate that the MC38 ICI response is controlled by multiple host genes, with epistatic interactions taking place between subsets of these genes.

### IMMUNE GENES WITHIN QTL ARE THE KEY DRIVERS OF ICI RESPONSE IN THE CCF1 MC38 SYSTEM

Our genetic mapping uncovered four QTL harboring genes whose variations in either structure or expression significantly impact on ICI response. Since the QTLs span large segments of the genome (approximately 5-50 megabases each), the total number of candidate quantitative trait genes (QTGs) is high (*N* = 2,793). We therefore conducted an integrative analysis of all four QTL to understand whether these genes showed differences between responder and non-responder strains, and what biological functions were enriched among these collective gene sets. First, we obtained bulk transcriptional profiles of MC38 tumors (*N* = 23) from three responder strains (CC01 F1, CC02 F1, CC75 F1) and three non-responder strains (CC36 F1, CC79 F1, CC80 F1) treated with either isotype or aPD1. Of 22,419 genes with detectable expression in these tumors (≥5 reads across samples), 2,395 (10.7%) were significantly differentially expressed genes (DEGs) between responder and non-responder tumors (FDR = 5%).

Given the fundamental importance of immunobiology to ICI response, we sought to understand the degree to which genes with known immune-related functions were reflected in our QTL and the DEGs within QTL. We leveraged the InnateDB resource^83^ to compile a systematically annotated list of genes (*N* = 8,400) associated with immune function. We found that genomic loci spanned by our QTL (regardless of gene expression) harbored a slightly higher fraction of immune-related genes compared to the genomic background (Fisher’s test *P* = 0.05). However, DEGs distinguishing responders from non-responders within our QTLs were very strongly and significantly enriched for immune function compared to non-DEGs within QTL (58% vs 31%; Fisher’s test *P* = 3.5×10^−11^).

We next explored the biological processes associated with immune vs. non-immune DEGs within our QTLs. For the immune DEGs within QTLs, top enriched KEGG pathways (all *P <* 1×10^−10^) included antigen processing and presentation, allograft rejection, and graft vs host disease (**Figure 2G**), which is consistent with the enhancement of responder strain anti-tumor (anti-self) immune responses induced by immune checkpoint inhibitors. Other top enriched KEGG pathways (all *P <* 1×10^−10^) included pathways related to viral infection (Epstein-Barr virus, myocarditis, Human T-cell leukemia virus) or autoimmune processes (Type 1 diabetes, thyroid disease; **Figure 2G**). These results likely reflect the role of CTL and interferon responses in immune regulation of the TIME^84^ and the balance that must be struck between the response against self-antigens and the destruction of malignant cells in a successful immune response to cancer^85^. In contrast to these strongly enriched pathways among immune DEGs within QTLs, there were no statistically significant enriched pathways among the non-immune DEGs within QTLs suggesting the absence of cohesive functions in these DEGs.

Among DEGs with immune function located within QTL, over half were derived from the mChr17 QTL (*N =* 51 [mChr17], 26 [mChr9], 12 [mChr15], and 4 [mChr5]). Except for the mChr17 QTL, none of the DEGs in the remaining individual QTLs associated strongly with a specific immune function (no enrichment of specific pathways with *P* < 0.001). The mChr17 QTL appears to be a special case since it encompasses the entire MHC locus, which includes MHC class I and II as well as immune genes such as TNF and lymphotoxin. The genes in this mChr17 QTL were strongly enriched for pathways representative of antigen presentation, autoimmunity, and interferon responses.

Since there are DEGs outside the QTLs that are immune related, we asked if there were quantitative and qualitative differences between those immune DEGs within the QTLs and those outside the QTLs. While we found that immune DEGs within all QTLs were strongly enriched for pathways linked to cellular immunity and immunotherapy responses (antigen presentation, autoimmunity, interferon responses, Th1/Th2/Th17 cell differentiation, and T cell receptor signaling), the immune DEGs outside the QTL were enriched for pathways that were not as obviously associated with anti-cancer immunity (e.g. response to bacterial infection, B cell receptor signaling, apoptosis). Taken together, these results suggest that there is functional coherence in the major host genetic drivers residing within our QTLs for ICI response and that these drivers interact with each other to establish a responsive TIME.

### IMMUNOPHENOTYPING REVEALS SPECIFIC CHARACTERISTICS ASSOCIATED WITH THE ICI-RESPONSIVE TUMOR IMMUNE MICROENVIRONMENT

Given the impact of QTL immune genes on ICI responsiveness, we sought to define the differences in the TIME between responder vs. non-responder CCF1 strain MC38 tumors. To accomplish this, we first examined changes in cellular composition within tumors. To this end, we selected the same three consistent responder strains (CC01 F1, CC02 F1 and CC75 F1) and three non-responder (CC36 F1, CC79 F1 and CC80 F1) strains for immunophenotyping based on their response in our initial screen (**Figure 1B**). Forty-eight hours after a single dose of either aPD1 or isotype control, when the tumors show no macroscopic changes in size between treatment groups, we harvested MC38 tumors and characterized their cell composition using both single cell RNA-Seq (scRNA-Seq) and flow cytometry (**Figure 3A**). This approach ensured that any changes we quantified were not due to volume changes in the tumors.

**Figure 3:**
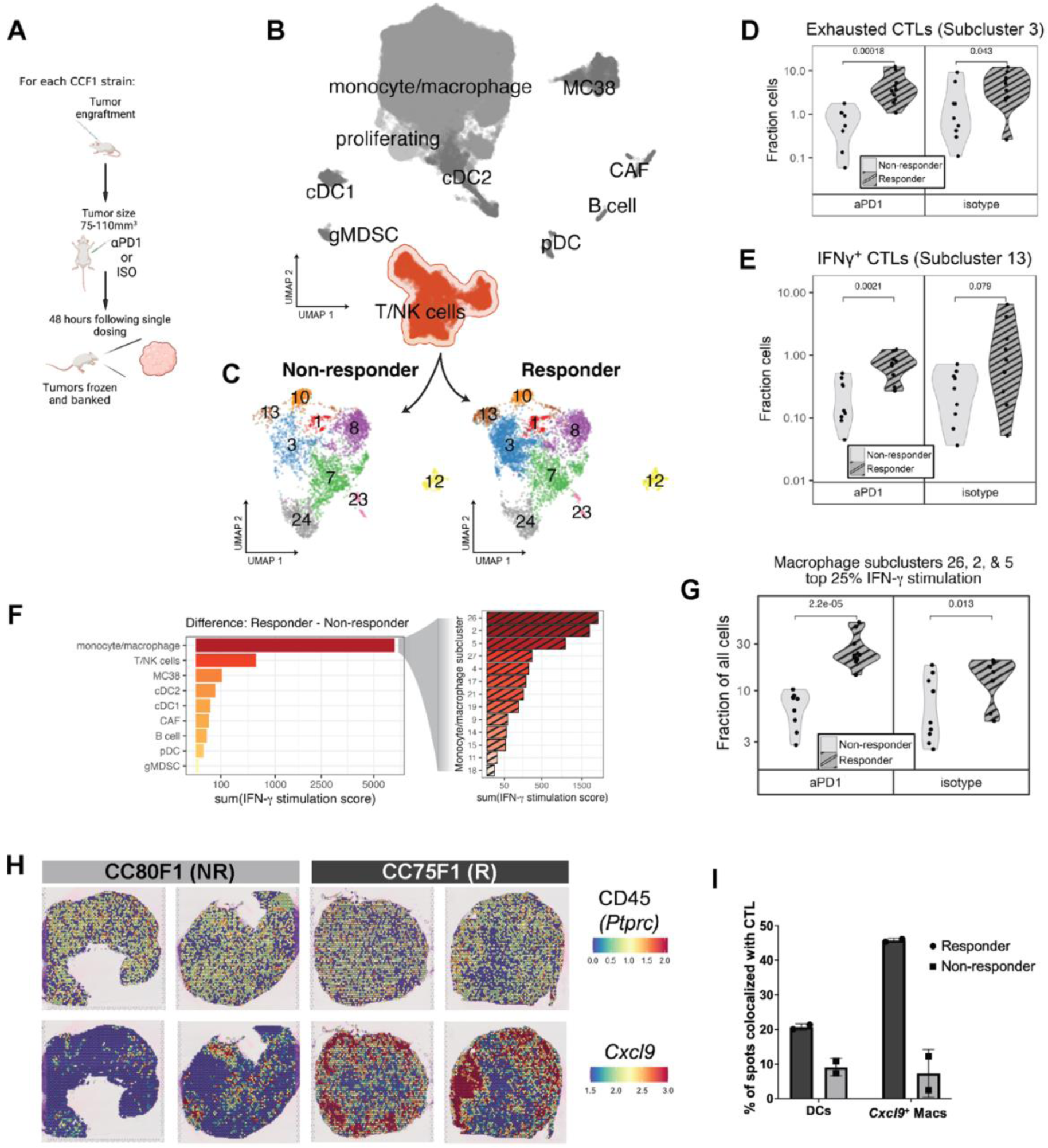
Immunophenotyping reveals enrichment and co-localization of CTL and IFNγ-stimulated macrophages in responder strain MC38 tumors. (A) Overview of single dose paradigm where tumors are harvested 48 hours after a single dose of aPD1 or isotype control in order to capture the initial effects of treatment prior to macroscopic changes in size. (B) UMAP plot depicting an overview of heterogeneity in single cell transcriptomes derived from responder and non-responder strain tumors and highlighting the T/NK cell cluster. (C) Subclustering of the T/NK cell cluster reveals transcriptionally distinct cell subsets showing enrichment of subclusters 3 and 13 in responding tumors. (D) Fraction of exhausted CTLs (cluster 3) as portion of all cells in responder and non-responder strain groups. (E) Fraction of IFNγ^+^ CTLs (cluster 13) as portion of all cells in responder and non-responder strain groups. (F) Expression of an 18-gene signature reflecting IFNγ^+^ stimulation applied to all cells in each cluster (left) or macrophage subcluster (right), with bars depicting the difference in summed pathway score between responder and non-responder cells within each (sub)cluster. (G) Fraction of IFNγ stimulated macrophage clusters (clusters 2, 5, and 26) as portion of all cells in responder and non-responder strain groups. (H) Visium spatial transcriptomics images showing expression of CD45 (*Ptprc*) [top row] and *Cxcl9* [bottom row] in spatially-resolved spots of tumors harvested from responder (CC75F1) and non-responder (CC80F1) mouse tumors. (I) Percent of spatial transcriptomics spots expressing markers indicative of DCs and *Cxcl9^+^* macrophages that are co-localized with expression of CTL markers.

Flow cytometry revealed no significant difference in the overall infiltration of CD45^+^ immune cells between tumors from responder (N = 28) and non-responder (N = 25) strain mice either in the isotype treated or in the single dose aPD1 treated groups (**Table S2**). Specifically, the immune cellular composition of the MC38 TIME was constant with CD11b^+^F4/80^+^ macrophages dominating at ∼70% of CD45^+^ cells, followed by F4/80^−^CD3e^+^ T cells (∼20% of CD45^+^ cells), and then dendritic cells (DCs; ∼10% of CD45^+^ cells) regardless of the treatment or the response group (**Table S2**). These proportions are similar to what had been previously described for the MC38 model^86^. Moreover, our data showed intratumor enrichment of MC38 antigen-specific MHC class I restricted tetramer-specific CTL and PDL1^high^MHC class II^high^ macrophages in MC38 tumors of responder CCF1 strains (**Table S2**).

To further dissect immune cell heterogeneity in MC38 tumors, we built a single cell transcriptomic atlas of MC38 tumors using scRNA-Seq. We used MC38 tumors from our single ICI dose protocol (**Figure 3A**) capturing a range of responder (*N* = 20; tumors from CC01 F1, CC02 F1, and CC75 F1) and non-responder (*N* = 18; tumors from CC36 F1, CC79 F1, and CC80 F1) strain tumors. Following sample acquisition, we profiled tumors using the 10X Genomics Chromium platform. We sorted to enrich for the CD45^+^ compartment (Methods) and recovered approximately 76,000 cells consisting of a mix of cells derived from responder and non-responder CCF1 strain tumors.

We used standard methods to cluster single cell transcriptional profiles and classified 9 major cell populations (**Figure 3B**). Again, we identified populations of monocyte/macrophages (*Adgre1, Apoe, Lyz2*) as the predominant tumor-associated immune component, followed by T/NK cells (*Cd3e, Nkg7, Ifng*), with B cells (*Cd79a, Igkc, Ms4a1*), conventional dendritic cells (cDC1: *Xcr1, Clec9a, Irf8*; cDC2 *Irf4, Itgam, Sirpa*) plasmacytoid dendritic cells (pDCs: *Klk1, Igkc, Siglech*), cancer associated fibroblasts (CAFs: *Col1a1, Bgn, Postn*), and granulocytic myeloid-derived suppressor cells (MDSCs: *Gsr, Retnlg, S100a9*) accounting for a significantly smaller proportion of the tumor-associated stroma. These were present in approximately the same proportion as was found in the flow cytometry data (**Table S2**).

To obtain a more refined portrait of differences between the responsive and non-responsive TIME, given the importance of the T cell response in anti-tumor immunity, we began by subclustering the heterogeneous T/NK cell cluster (**Figure 3C**). We assessed cell composition and gene expression differences between responder and non-responder tumors. We found two CTL subclusters that were significantly enriched in responder strain MC38 tumors (**Table S2 and Figure 3D-E**). Differential gene expression analysis of these CTL subgroups (scRNA-Seq subclusters 3 and 13) revealed a cassette of specific marker genes that identified CTLs (*Cd3e, Cd8a, Cd8b1*), and either markers of exhaustion (subcluster 3; *Tox*, *Lag3, and Pdcd1* [PD-1])^87^ (**Supplemental File Tab A**), or cytotoxicity (subcluster 13; *Ifng, Gzmb, Prf and Tbx21*) (**Supplemental File Tab B**). The cytotoxicity subcluster (subcluster 13) exhibited the highest expression of *Ifng* (IFNγ) when compared to other cells in the dataset (>34 fold) (**Supplemental File Tab B**)^88^. The proportions of these CTL clusters were higher in responder tumors regardless of treatment (either isotype or aPD1, **Figure 3D-E**). Thus, enrichment of *Tox*^+^ exhausted-like and *Ifng*-expressing CTLs defining the responder strain TIME were established prior to ICI exposure rather than as a response to aPD1 therapy.

Because *Ifng*^+^ CTL were enriched in responder strain tumors, we examined which cells might be stimulated by IFNγ in responder strain tumors. To assess stimulation by IFNγ, we used a previously published 18 gene IFNγ-related mRNA profile (IFNγ response signature, “Merck18”) that predicts clinical response to PD-1 blockade^89^. We compared cells using this response signature and found that macrophages showed a prominent difference between responder and non-responder samples (**Figure 3F, left**). We then subclustered macrophages and asked whether the proportion of cells that scored as highly responding to IFNγ (upper quartile score) in each macrophage/monocyte subcluster differed between responder and non-responder strain tumors. Of the ten total monocyte/macrophage subclusters identified in the scRNA-seq dataset (methods), three subclusters – 2, 5, and 26 – contained the bulk (82%) of macrophages with high IFNγ response signature scores (**Figure 3F, right**), suggesting that these macrophage subpopulations were primarily responding to IFNγ stimulation. These three subpopulations comprised a substantial fraction of our complete dataset, with macrophage subclusters 2, 5 and 26 representing 20%, 18%, and 13% of all cells collected, respectively. Cells within macrophage subclusters 2, 5 and 26 that had high IFNγ response signature scores (upper quartile) were significantly enriched in responder versus non-responder strain tumors (**Table S2, Figure 3G**). PD-L1 and MHC class II are IFNγ-inducible genes^89^, and our FACS data using these markers confirmed that a PD-L1^high^MHC class II^high^ M1-like macrophage population^90^ was enriched in the responder strain TIME (**Table S2**).

These data suggested a potential cross-talk between CTLs and macrophages. We therefore asked whether these interactions are facilitated by co-localization of these cell types within tumors. To test this, we used the 10X Genomics Visium spatial transcriptomics to ask if the IFNγ-stimulated (*Cxcl9*^+^) macrophage population^89^ and CTLs were distributed differently between tumors from responder (CC075F1, *N* = 2) vs. non-responder strains (CC080F1, *N* = 2) two days after treatment with a single dose of isotype antibodies to represent a pre-aPD1 treated tumor TIME state. Analysis of the expression of *Ptprc* (CD45), the pan-immune cell marker, suggested that both responder and non-responder tumors were similarly diffusely infiltrated by immune cells (**Figure 3H**). In contrast, examining the IFNγ responsive marker *Cxcl9*^89^, responder tumors displayed substantially more positive spots with clear punctate foci, suggesting intratumor clusters of *Cxcl9*^+^ monocyte/macrophages (**Figure 3H**). An examination of the CTL marker, *Cd8a,* revealed a similar pattern to *Cxcl9* (data not shown). By computing the fraction of *Ptprc*^+^ spots co-expressing either macrophage markers (*Adgre1, Itgam,* and *Cxcl9*) or DC markers (*Batf3* and/or *Zbtb46*) and expressing the CTL markers *Thy1* and *Cd8a*, we found that responder strain tumors showed prominent co-localization of *Cxcl9*^+^ macrophages with CTLs: responder strain tumors have >4X CTL-macrophage co-localized spots and 2X more DC-CTL interactions than non-responder strain tumors in this pre-treated state (**Figure 3I**).

### ANALYSIS OF RESPONSE ASSOCIATED GENES WITHIN THE MURINE QTL UNCOVERS A TOP-PERFORMING TRANSCRIPTIONAL PREDICTOR OF SURVIVAL IN SPECIFIC HUMAN CANCER IMMUNE SUBTYPES

After establishing the importance of immune genes in our QTL, and the differences in TIME between responders and non-responders, we sought to identify key QTGs mediating variation in ICI response in our system. We therefore developed an integrative strategy that prioritizes candidates most likely to strongly impact ICI response across species. This approach combines differential gene expression within the tumor microenvironments of responder and non-responder mice along with cross-species validation in large human cohorts. We hypothesized that these genes instantiate the fundamental biology of TIME composition, and that variations in the expression of these genes may reflect the degree to which a particular TIME is primed for responsivity to ICI.

A recent study of human patients identified macrophage polarity defined by the expression ratio of the genes *CXCL9* and *SPP1* (but not by conventional M1 and M2 markers) as having a strong prognostic association with survival in several cancers, as well as associating with response to aPD1^53^. To test whether our MC38 system recapitulated this association in humans, we examined bulk transcriptomes of 23 MC38 tumors from responder and non-responder strain CCF1 mice treated with aPD1 or isotype control using the single dose protocol previously described. This approach assesses transcriptional dynamics prior to any observable ICI-induced effects on tumor size. An examination of *Cxcl9* and *Spp1* expression in these tumors revealed that the MC38 tumors in responder CCF1 strains treated with isotype control antibody showed a significantly greater *Cxcl9*/*Spp1* ratio than those from non-responder strain mice (**Figure 4A**). Intriguingly, a single dose of aPD1 treatment further augmented the *Cxcl9*/*Spp1* ratio in responder strain tumors by >2-fold whereas there was no change in this ratio in non-responder strain tumors. This resulted in a much greater difference in the *Cxcl9*/*Spp1* ratio in responder strain MC38 tumors after treatment than non-responder strain tumors (**Figure 4A**). These results demonstrate that the MC38 CCF1 system is not only a cross-species validation of the predictive value of the reported *Cxcl9*/*Spp1* ratio for ICI anti-tumor response, but also suggests that a TIME propitious for ICI response in our CCF1 system is reflective of the human condition.

**Figure 4:**
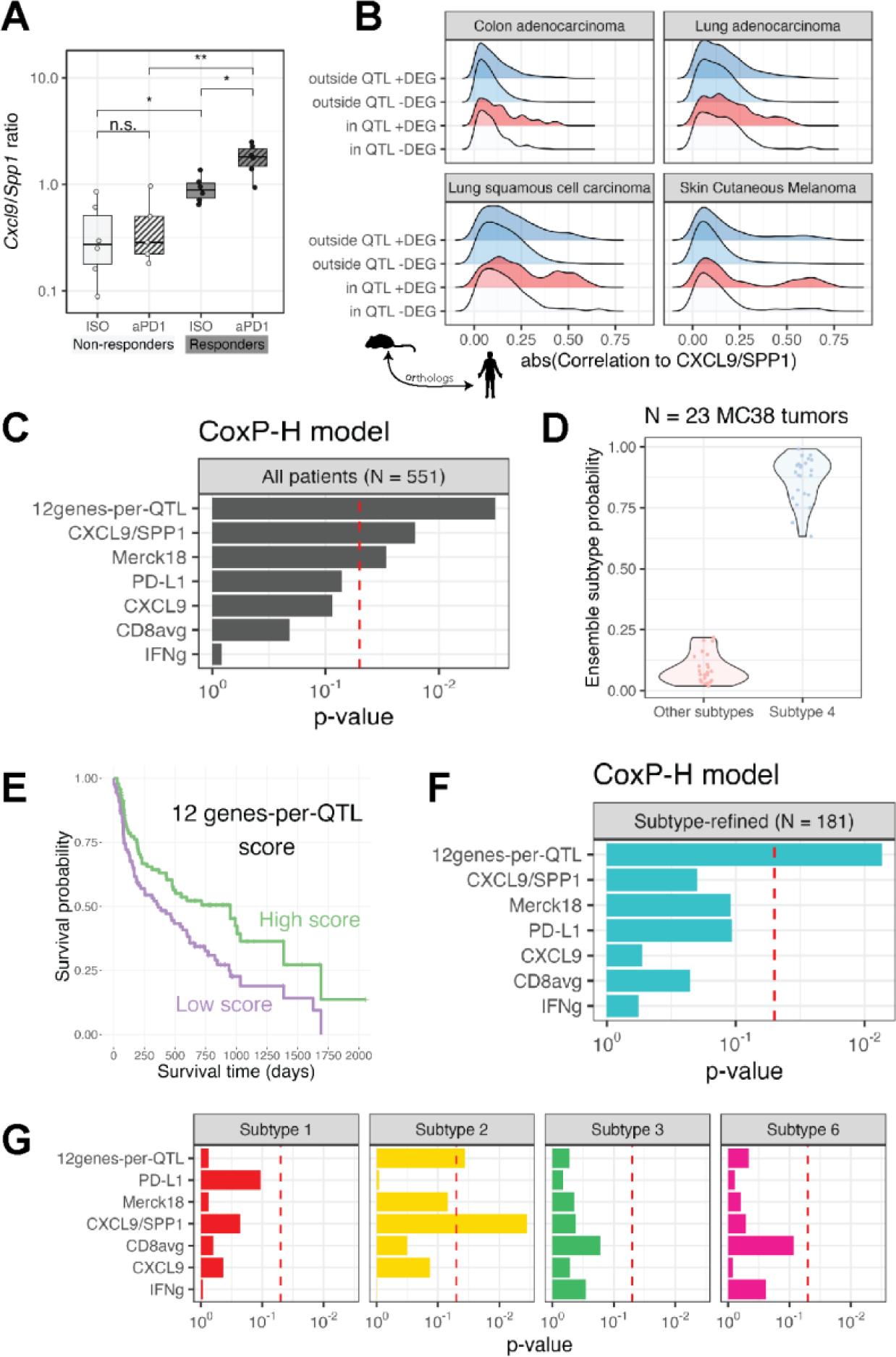
CST algorithm prioritizes genes within murine QTL predicting survival in specific human cancer immune subtypes. (A) *Cxcl9/Spp1* ratio in MC38 bulk RNA-Seq samples from responder and non-responder strain tumors. \*\**P* < 0.01; \**P* < 0.05; n.s. – not significant (B) Ridge plot depicting the distribution of correlation coefficients between each human homologue of the murine genes with the *CXCL9/SPP1* ratio in four TCGA cohort. Gene sets are divided based upon whether each gene is located within or outside our four murine QTL and whether the gene was a DEG or not in bulk RNA-Seq of responder and non-responder strains. The only gene set comprising of genes that are within the QTL and a DEG show a subset highly correlated with *CXCL9/SPP1* ratio in human tumors. (C) Cox proportional hazards analysis showing the predictive significance of transcriptional biomarkers derived using the various genesets shown. (D) The predicted tumor immune subtype of *N* = 23 MC38 tumors is subtype 4. (E) Superior survival is seen in patients undergoing aPD1 treatment who are high expressors (defined as having a score above the median) of our 12 genes-per-QTL transcriptional biomarker (F) Cox proportional hazards analysis showing the predictive significance of the 12 gene-per-QTL transcriptional biomarkers over all other biomarkers when tested on only patients with immune subtype 4 tumors treated with aPD1 therapy. (G) As in (F), but transcriptional biomarkers were applied to patients with immune subtypes 1, 2, 3, or 6 as shown in facets.

With the knowledge that the *CXCL9/SPP1* ratio is associated with human cancer outcomes and highly predictive of response in our MC38 system, we leveraged The Cancer Genome Atlas (TCGA) data^52^ as a filter for cross-species translation. We focused on cancers that were similar in tissue origin to our MC38 model (colon) or are commonly treated with aPD1 immunotherapy (lung, melanoma), resulting in four TCGA cohorts (*N* > 500 for all cohorts). In these cohorts, we used the *CXCL9/SPP1* ratio as a proxy for a favorable TIME that is likely to respond to ICI. We directly computed the correlation of the human ortholog of each mouse gene to *CXCL9/SPP1*. As expected, we found a range of correlations to this ratio, with most genes having a low absolute correlation magnitude (<0.25, Spearman rank correlation coefficient). However, density plots revealed that the magnitude of correlation to *CXCL9/SPP1* of DEGs within our QTL had a distinct “long tail” of genes with much higher correlations to this ratio (**Figure 4B**, red). In contrast, non-DEGs outside QTL did not present a meaningful fraction of genes falling within this long tail. We surmised that the long tail of DEGs with elevated correlations to *CXCL9/SPP1* ratio within our QTL represent the critical QTGs – genes that are instrumental in the priming of a favorable TIME that enables response to ICI in a cross-species manner. We used the distribution of *CXCL9/SPP1* correlations of non-DEGs outside QTL as an empirical null distribution to compute the significance of those correlations of DEGs within QTL. We refer to this approach as the cross-species TIME (CST) algorithm. In this manner we were able to utilize the CST algorithm to rank genes within QTL discovered in our mouse system according to a measure of their influence on the TIME in human patients.

Using this ranking approach, we identified the human homologs of the top 100 genes based on the CST ranking within the four murine QTL associated with ICI response in our MC38 system (Table 1). To explore whether these genes have prognostic value in human ICI patients, we assembled a cohort of *N =* 551 patients treated with aPD1 therapeutics and with pre-treatment tumor transcriptional profiles (ICI validation set, *N* = 7 different studies and cancer types^54,55^) and tested different genesets on their performance in predicting aPD1 response. To formally test the prognostic value of our score on overall survival, we used a stratified Cox proportional hazards model. For each geneset, we computed a single composite score that equally weighted the contributions of each gene (Methods). Because we demonstrated epistatic interactions between specific QTL, we assembled genesets including genes from each of our QTL. We found best survival prediction when using 12-18 genes per QTL **(Figure S2A**). A minimum geneset of 12 genes-per-QTL (total 48 genes) significantly predicted overall survival in the 551 patient cohort (*P* = 0.0028). We compared our score to other commonly used transcriptional biomarkers of ICI response and found that it outperformed all other metrics in predicting overall survival, although two other biomarkers – *CXCL9/SPP1* ratio^53^ and Merck18^89^ – were also significantly predictive (*P <* 0.05; **Figure 4C**). When we compared this 48 pan-QTL geneset with a set of 48 genes selected strictly on CST rank while disregarding QTL assignment, we found that, while both genesets were predictive, the pan-QTL geneset outperformed the top CST-ranked 48 geneset (*P =* 0.0028 vs. *P* = 0.024, respectively), consistent with the importance of epistatic interactions across the QTLs associated with the responsive TIME.

**Table 1:**
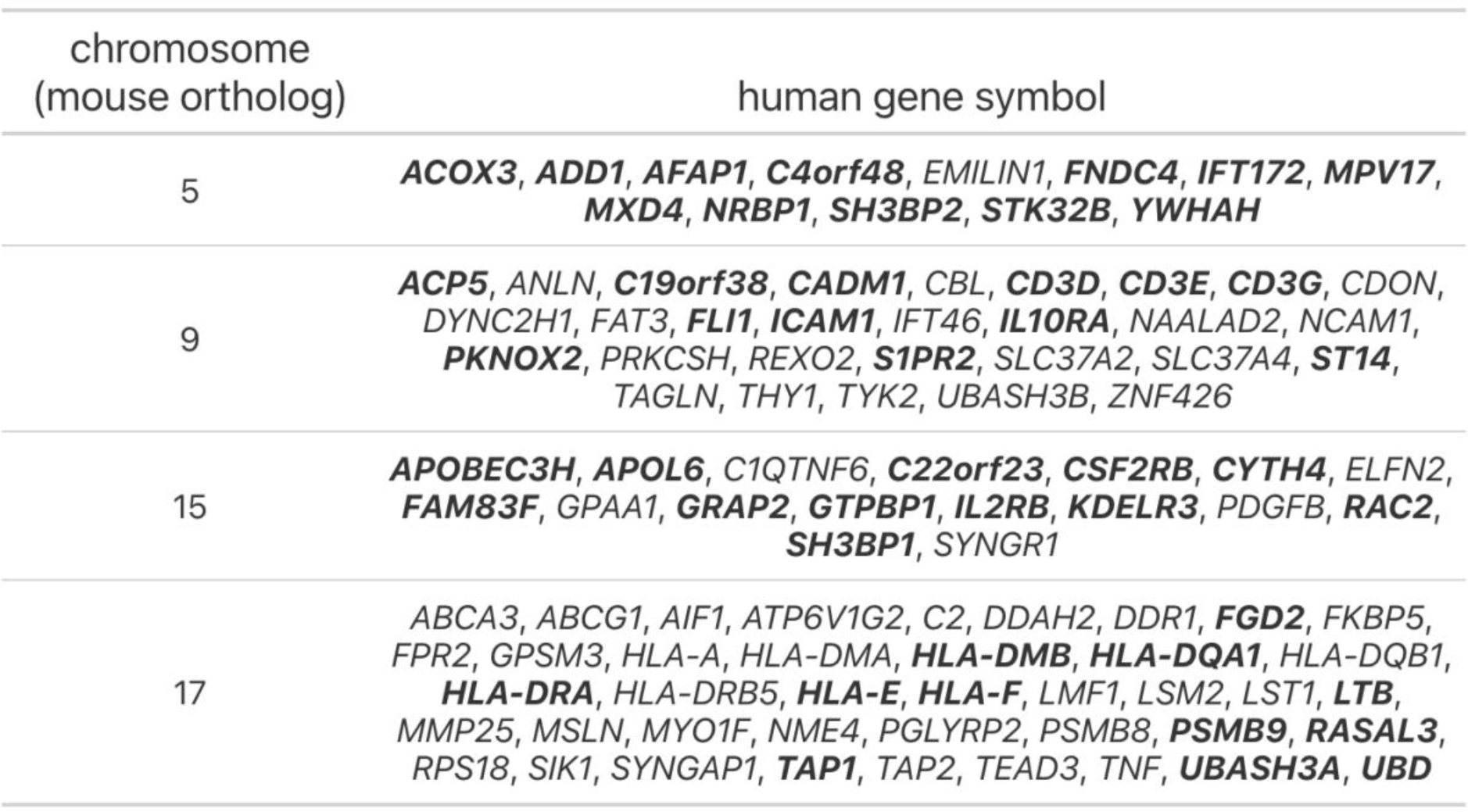
Candidate genes prioritized using our CST algorithm predicted to have a major role establishing a tumor immune microenvironment favorable for aPD1 response in mice and humans. Candidates in bold are the top twelve for each QTL that comprise the 48-gene panel used as the predictive markers in Figure 4.

| chromosome<br>(mouse ortholog) | human gene symbol |
| --- | --- |
| 5 | **ACOX3, ADD1, AFAP1, C4orf48, EMILIN1, FNDC4, IFT172, MPV17, MXD4, NRBP1, SH3BP2, STK32B, YWHAH** |
| 9 | **ACP5, ANLN, C19orf38, CADM1, CBL, CD3D, CD3E, CD3G, CDON, DYNC2H1, FAT3, FLI1, ICAM1, IFT46, IL10RA, NAALAD2, NCAM1, PKNOX2, PRKCSH, REXO2, S1PR2, SLC37A2, SLC37A4, ST14, TAGLN, THY1, TYK2, UBASH3B, ZNF426** |
| 15 | **APOBEC3H, APOL6, C1QTNF6, C22orf23, CSF2RB, CYTH4, ELFN2, FAM83F, GPAA1, GRAP2, GTPBP1, IL2RB, KDELR3, PDGFB, RAC2, SH3BP1, SYNGR1** |
| 17 | **ABCA3, ABCG1, AIF1, ATP6V1G2, C2, DDAH2, DDR1, FGD2, FKBP5, FPR2, GPSM3, HLA-A, HLA-DMA, HLA-DMB, HLA-DQA1, HLA-DQB1, HLA-DRA, HLA-DRB5, HLA-E, HLA-F, LMF1, LSM2, LST1, LTB, MMP25, MSLN, MYO1F, NME4, PGLYRP2, PSMB8, PSMB9, RASAL3, RPS18, SIK1, SYNGAP1, TAP1, TAP2, TEAD3, TNF, UBASH3A, UBD** |

While our 12genes-per-QTL composite score has prognostic value for survival in ICI-treated patients, it may not be equally informative across all types of tumors. In a previous study of TIME characteristics, Sayaman et al.^32^ showed that trait heritabilities varied in part based on which of six previously defined tumor immune subtypes^11^ each tumor matched. We leveraged a machine learning method to predict immune subtypes in our mouse MC38 RNA-Seq samples^65^ (Methods). We found that all MC38 tumors (N=23), regardless of strain of origin (3 responder and 3 non-responder strains) or treatment group (isotype vs aPD1), were classified as immune subtype 4 (**Figure 4D**), which though originally described as “lymphocyte depleted” is characterized by a high macrophage:lymphocyte ratio^11^. We used the same method to assign immune subtypes of the 551 patients in our ICI validation set and found that 181 tumors were classified as immune subtype 4. Bifurcating patients into high vs low-scoring based upon cohort-specific thresholds, we observed significantly longer survival (*P* = 0.0054, log-rank test) in subtype 4 patients with a high 12genes-per-QTL composite score (**Figure 4E**). Using the proportional hazards approach on this subtype-refined set of 181 patients, our composite score was significantly predictive of overall survival (*P* = 0.0092), while no other biomarker, including the *CXCL9/SPP1* ratio used in the CST algorithm, retained predictive capacity (**Figure 4F**). Among tumors of other specific immune subtypes present in our ICI validation set, our score was also marginally predictive for subtype 2, for which the *CXCL9/SPP1* ratio performed best (**Figure 4G**). These results reinforce that the QTGs identified using our mouse system are not general indicators of prognosis but specifically reflect fundamental and conserved TIME biology of tumors matching specific tumor immune subtypes.

The importance of the MHC locus is underscored not only by how the mChr17 QTL is significantly engaged in epistatic interactions with the QTLs on mChr 5 and 15, but also that nine of the top 100 (and five of the 48 pan-QTL geneset; **Table 1**) response-associated DEGs were MHC genes in responsive tumors. The human homologues of these top genes identified by the CST algorithm were both HLA class I (HLA-A, HLA-E, HLA-F) and class II genes (HLA-DMA, HLA-DMB, HLA-DQA1, HLA-DQB1, HLA-DRA, HLA-DRB5). Mice that are homozygous for shared B6 and 129 haplotypes (H2^b^ including H2-K^b^) in mChr17 exhibit the best responses as 129 is haploidentical (including H2-K^b^) with B6 at the MHC locus. H2-K^b^-restricted tetramer analysis on responder vs non responder tumors after a single dose of aPD1 revealed that responders showed a significantly greater frequency of H2-K^b^-restricted MC38-specific (against the p15E antigen) CTL response than non-responder tumors (∼10 fold; **Figure S2B**). These data suggest that B6/129 homozygosity in this specific context augments MC38-specific CTL responses over all mice heterozygous at mChr17 likely through more efficient presentation of the MC38 specific, H2-K^b^-restricted antigen. As this identical biallelic configuration is a unique circumstance caused by controlled breeding of inbred mouse strains, we examined the relationship between MHC variability and aPD1 response in the other CCF1 lines (**Figure S2C**). We eliminated the NOD strain from this analysis since it is known to have deficits in the maturation and function of antigen-presenting cells^91^. Using ∼84,000 SNPs in the murine MHC region genotyped in the eight inbred CC founder strains, we computed the heterozygosity of the MHC locus of each CCF1 cross based on the level of SNP diversity at this locus. We found that increasing heterozygosity at the MHC locus is associated with improved ICI response as measured by our RTC metric (**Figure S2D**).

### IN VIVO BLOCKADE OF GM-CSF AND IL2RB REVERSES THE TRANSCRIPTIONAL SIGNATURE OF aPD1 RESPONSE

Using our CCF1/CCF1N1 approach we have uncovered highly plausible candidate drivers that shape an ICI-favorable TIME in both mice and humans. These top response-associated genes were not expressed focally in any one immune cell type, and did not group within a single pathway, suggesting that broad cellular and genetic networks are responsible. This raised the question of whether simultaneous perturbation of a number of these QTGs is required before the aPD1 response can be altered. To address this question, we examined two top candidate QTGs on mChr15 in our 48 gene panel for which there are validated murine reagents, *Csf2rb* (GM-CSF receptor), and the *Il2rb* (IL2 receptor beta subunit). The addition of their cognate cytokines, GM-CSF and IL2 (i.e., IL2RB-targeted pegylated form), have been noted to enhance ICI response in both preclinical and some clinical studies^92,93^. In examining the single cell genomics data of our MC38 tumors, we found that the genes encoding the three subunits comprising both the high-affinity IL-2 receptor (*Il2ra, Il2rb, Il2rg*) and the IL-15 receptor (*Il15ra, Il2rb, Il2rg*) were co-expressed in a minority of cells but these were significantly enriched in CTL clusters (binomial test, P < 1×10^−10^) suggesting that the functional receptor complexes are present in the appropriate immune cell populations in our system. Therefore, our genetic analyses would predict that blockade of the receptors or the cytokines would attenuate aPD1 response in a responder genetic background. To test this prediction, we chose a responder CCF1 cross – CC75×B6. We have previously determined that a single dose of aPD1 can dramatically augment the *Cxcl9/Spp1* ratio in responder mice, and act as a surrogate biomarker of aPD1 response. With this framework, we first applied a GM-CSF blocking antibody to MC38 tumor-bearing responder mice using standard protocols^94^ (**Figure 5A**) and assessed the *Cxcl9/Spp1* ratio 48 hours after administration. While the addition of aPD1 significantly augmented the *Cxcl9/Spp1* ratio in MC38 tumors, the co-administration of aPD1 + anti-GM-CSF (aGM-CSF) completely abrogated the induction of the ratio to baseline levels (**Figure 5B**). When we directly blocked the IL2RB receptor using anti-IL2RB antibody (aIL2RB)^95^, the *Cxcl9/Spp1* ratio was reduced even further to levels similar to those found in tumors of genetic non-responders (**Figure 5B, 5F**) and intratumor CTL infiltration appeared to be compromised (**Figure S3, Figure 5F**). When the two blocking antibodies were administered together with aPD1, the ratio declined even further but this difference was not statistically significant.

**Figure 5:**
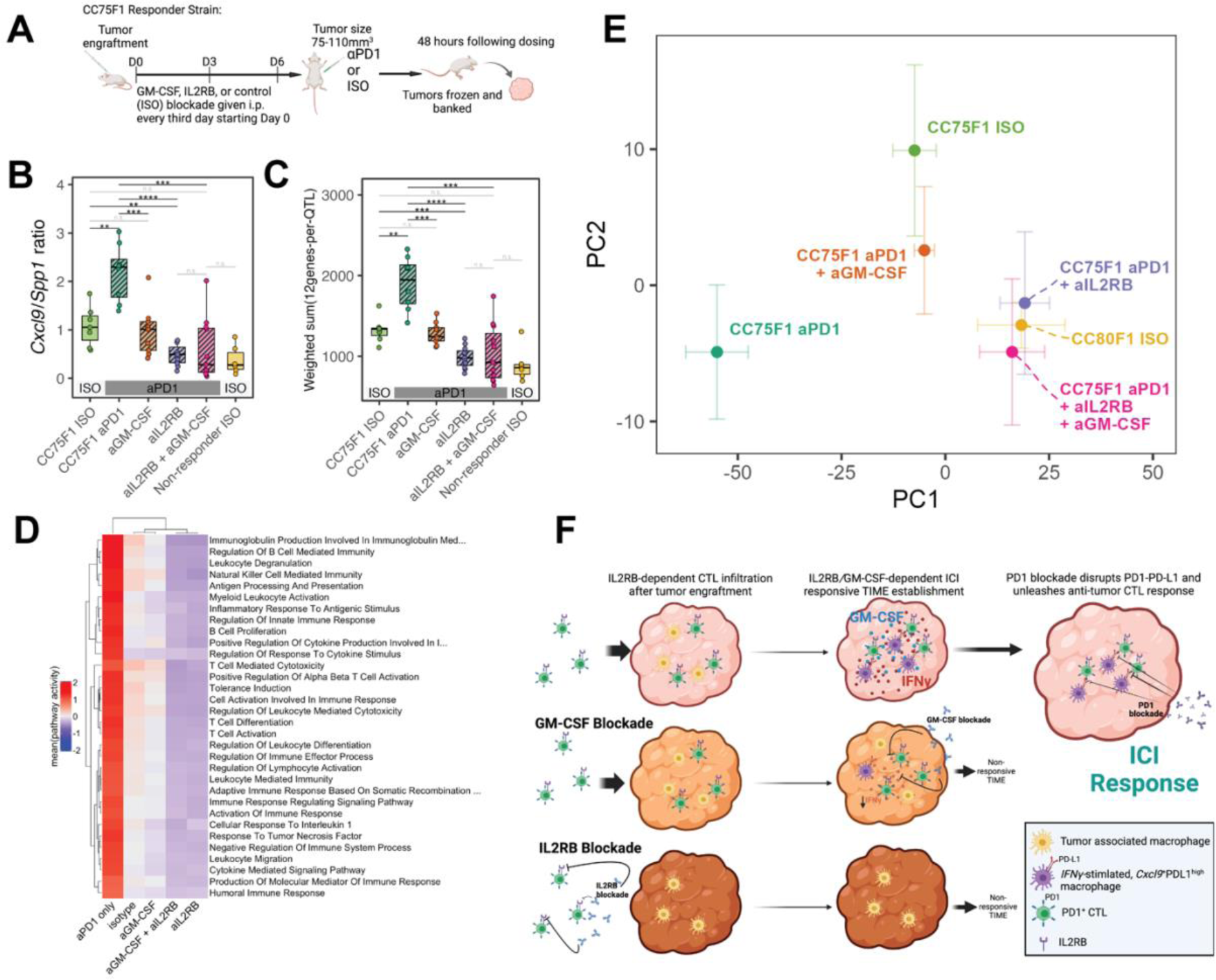
*In vivo* blockade of GM-CSF and IL2RB reverses the transcriptional signature of aPD1 response. (A) Schematic overview of the blocking antibody dosing protocol and strategy for assessing tumors before macroscopic changes in size. (B) *Cxcl9/Spp1* ratio computed using RNA-Seq gene expression profiling of CC75F1 MC38 tumors collected as in (A). (C) Transcriptional changes of our 48 TIME gene panel assessed on the same tumors as in (B). \*\*\*\**P* < 0.0001; \*\*\**P* < 0.001; \*\**P* < 0.01; \**P* < 0.05; n.s. – not significant (D) Heatmap of enriched immune-related pathways among the experimental groups profiled in this blocking antibody experiment showing reversal of immune signatures. Pathway activity was computed based on leading edge genes identified by gene set enrichment analysis (Methods) and constituted the mean of standardized (per-gene, across all samples) gene expression. (E) Principal component (PC) plot showing the positioning of each group of samples within the first two PCs. CC75 ISO is the tumor in a responding genetic cross treated with isotype control antibodies whereas CC80 ISO is the tumor in a non-responding genetic cross treated similarly. Principal component analysis was performed using batch-corrected RNA-Seq profiles (Methods) with error bars showing +/−1 standard error within each group. (F) Model describing the effects of IL2RB and GM-CSF blockade on the MC38 TIME and response to aPD1.

To confirm that these changes were not restricted only to the two transcripts (*Cxcl9* and *Spp1*), we assessed the transcriptional changes of our 48 pan-QTL gene panel and found markedly similar patterns: aGM-CSF abrogated the aPD1 transcriptional response and aIL2RB significantly reduced that ratio further, while the combination of the two blocking antibodies did not significantly further reduce the ratio (**Figure 5C**). We explored pathway enrichment of genes differentially expressed in pairwise comparisons between any of the treatment groups (isotype control, aPD1 only, aPD1 + aGM-CSF, aPD1 + aIL2RB, aPD1 + both blocking antibodies). Among the immune-related enriched pathways, all were most highly expressed in the mice administered aPD1 only, with intermediate expression of these pathways in the mice administered isotype control or aGM-CSF + aPD1, and lowest expression in mice where IL2RB receptor was blocked (**Figure 5D**), following the rank-ordering observed for the *Cxcl9/Spp1* ratio (**Figure 5B**). Using a whole transcriptome approach and assessing the changes post treatment using principal component analyses, we found that, indeed, the co-administration of aGM-CSF with aPD1 returned the transcriptional configuration in CC75×B6 tumors to the responder baseline (i.e., CC75×B6 treated with isotype control), while aIL2RB treatment transformed the aPD1 induced transcriptome to the configuration of a genetic non-responder (CC80×B6 treated with isotype control; **Figure 5E, 5F**). These data not only validate our genetic discovery platform but point to an important biological possibility: while our QTL/QTG analysis implicated dozens of genes shaping the ICI response propitious TIME, perturbation of a single gene in this network is capable of completely disrupting the response transcriptional outcome.

## Discussion

The major limiting problem in assessing ICI response genetics is the complexity of the clinical patient population under study: every patient’s immune system is different because of host genetic variations; each tumor has unique genomic profile shaped by its own mutational origins and evolutionary path; each patient’s immune status has been further modulated by environmental factors, the microbiome, age, and prior anti-cancer treatment. These multiple independent variables preclude common approaches to unbiased and genome-wide association mapping in patients.

Murine systems to study host genetics of ICI response have, until recently, also been lacking due to the absence of genetic diversity^70^. In a recent study, however, Hackett, *et al*.^96^ used an approach to identify a locus on mChr13 that modulated response of the transplantable B16 melanoma cell line to combined aPD1 and anti-CTLA4 (aCTLA4) treatment. In their case, they crossed B6 with a panel from the Diversity Outbred (DO) resource and used 142 F1 crosses as their mapping population. Pursuing a candidate gene approach at this mChr13 locus, they pointed to the prolactin family as a candidate gene family. Although this elegant study highlighted the role of host genetics in ICI response, the outbred nature of the DO resource does not allow detailed gene-based functional interrogation and validation within an appropriate genetic context.

We have resolved this experimental problem by crossing mouse lines with defined genetic variability, the Collaborative Cross (CC) strains^74–76^, with the inbred strain specific to the murine tumor models for ICI response. The limited number of available CC strains precludes fine mapping yet can still nominate QTL while revealing outlier responder/non-responder strains that can be crossed for more powerful CCF1N1 mapping. These genetic maneuvers, which form the basis of our CCF1/CCF1N1 platform, overcome allogeneic rejection while introducing sufficient genetic heterogeneity for mapping studies. Importantly, this system significantly reduces experimental variables by restricting the analysis to single tumor cell lines grown to a consistent size *in vivo*, given a single drug protocol (aPD1), with animals in living conditions engineered to minimize the impact of diet and housing as confounding factors. In this manner, host genetic diversity becomes the key variable effecting ICI response. Importantly, biological findings can be replicated and further examined using the same replenishable CCF1 crosses, something that cannot be accomplished with the DO mapping approach.

Using this CCF1/CCF1N1 strategy, we show that the broad-sense heritabilities of ICI response were all statistically significant ranging from 18-40% across the four tumor cell lines used covering two distinct cancers (colon and breast) derived from two different genetic backgrounds (B6 and BALB). While it is difficult to estimate the clinical impact based on *H*^2^ statistics, in the MC38 model that gave the highest *H*^2^ (0.40), we saw the complete range of aPD1 response, from no response to complete response only by adjusting the host genetic background. The levels of heritability we observed for ICI response fall broadly within a range that has been observed for many other human quantitative traits. For example, heritability of coronary heart disease was estimated to be 38% for females and 57% for males^97^, and 37% for major depressive disorders^98^. While it is known that uncontrolled environmental factors (e.g. age as an “exposure”) will significantly vary the apparent heritability of a trait^99^, our system allowed us to control for obvious environmental factors (e.g. age, diet, housing). Despite this, we found significant variance in *H*^2^ across the tumor model systems used. The CCF1 crosses showing the best and the worst aPD1 outcomes largely differed between the different tumor models (see Figure 1G). This suggests that either different response QTL, different genes within our characterized QTL, or different epistatic QTL interactions define the optimal response for each tumor model. Moreover, the tumor-host interaction factors that may influence heritability remain unclear as there was no discernable systematic association between tumor genomic configuration and *H*^2^. The two cell lines giving the highest heritability (33-40%) had different tumor mutational burdens, are from different cancer types, and derived from different genetic backgrounds. Conversely, the two tumor models with the highest TMB, MC38 and CT26, had markedly different *H*^2^ (40% vs 19% respectively). Thus, there may be yet-to-be defined genetic or biochemical configurations of a tumor that require different host genetics to establish the optimum TIME for checkpoint inhibitor response. While this has been hypothesized, our system is uniquely structured to systematically address this possibility.

Two major challenges of genome wide association studies are the very large cohorts needed to resolve QTGs within QTLs and the difficulty in finding appropriate models or comparable clinical designs with which to biologically validate genetic findings. Our CCF1N1 system addresses both problems by reducing the need for highly refined mapping because of the nature of the CC resource, the unprecedented access to critical tissues in a timely fashion, and the ability to experimentally test the impact of any candidate gene in the same biological system (i.e., same genetic background).

While pairwise epistasis between markers in human studies can be achieved using standard GWAS approaches, the statistical requirements to detect such interactions are daunting. Using specific intercrosses designed from our original CCF1 screen, we were able to uncover epistatic interactions between QTL on mChr15 and mChr17, and between mChr17 and mChr5 in a highly efficient manner (number of mice, *N* < 300). That mChr17 appears to be a genetic “organizer” through epistatic interactions with both mChr5 and 15 is not unexpected: mChr17 harbored the greatest number of candidate QTGs of any of the QTLs, and also contains the murine MHC locus which is responsible for the antigen processing and presentation processes that is core to many relevant cellular immune functions. As expected, by removing the mChr17 QTL effect through a selective cross between CC12 and CC80 with matched ancestry at the MHC locus, the mChr17 QTL was eliminated, but a new QTL on mChr9 emerged.

A major question of any animal model system is how relevant it is to the human condition under study. We addressed this as it relates to our model in two ways. First, we found that the IFNγ-stimulated TIME associated with ICI response in the CCF1 crosses mirrored that seen in human tumors responding to ICI treatment^100^. Similar to our findings demonstrating enrichment and close proximities of CTL and *Cxcl9*^+^*Cd274*^+^ (PD-L1) macrophages in responder strain tumors, patients with NSCLC or metastatic colorectal cancer tumors responding to PD1 pathway blockade have higher numbers and spatial proximities of CD8^+^ and PD-L1^+^ cells as quantified by the Immunoscore-IC score^101,102^. Moreover, we found that responder strain tumors showed a distinct component shift to *Ifng* or *Pdcd1*(PD1)-expressing CTLs and IFNγ−activated *Cxcl9-* and PD-L1-expressing macrophage subsets, reflecting core immunological aspects of the response-propitious TIME also previously described in patients^103^. Our work, however, extends these findings by strongly suggesting that a large portion of the ICI response-promoting TIME is under host genetic control.

Our access to tumor tissues shortly after ICI treatment in a highly controlled manner allowed us to integrate transcriptional signatures from the same mapping subjects to dramatically narrow down the critical genes within the QTL. We devised a computational approach that leverages cross species impact which also addresses the concerns of the relevance of a murine system to human ICI response. Using our CST algorithm, we could rank order human homologs of our murine quantitative trait genes into those likely to be relevant to patients undergoing ICI treatment thus reducing 2,793 potential candidates to less than 100. We identified 48 primarily immune genes that had significant prognostic value for overall survival in human aPD1 trials that outperformed other known biomarkers of response. Strikingly, this 48 gene prognostic panel was the only biomarker predictive of responses to aPD1 therapeutics in immune subtype 4 human tumors, a subtype with poor prognosis^11^ which was the same immune subtype of our MC38 tumors. By contrast, the *CXCL9/SPP1* ratio outperformed our 48 gene panel in human immune subtype 2 tumors. These results again support the notion that there are different optimal host genetics that function in concert with specific tumor immune subtypes.

The *in vivo* functional blockade of two top candidate QTG pathways, GM-CSF (*Csf2rb*) and IL2/IL15 (*Il2rb*) resulting in the complete reversal of the aPD1 transcriptional response validates our mapping system and the associated CST algorithm as a relevant discovery platform. Intriguingly, our data suggest that the IL2RB blockade that affects the lymphocyte compartment is dominant over myeloid compartment driven by GMCSF blockade in immune endpoint regulation. Translationally, these two targets have already moved into the clinic. Based on positive preclinical studies (reviewed in ^104,105^), the cognate cytokines of those receptors, GM-CSF and IL2, have moved to clinical trials. The success of these studies led to the ECOG clinical trial (E1608) comparing aCTLA4 (ipilimumab) vs. recombinant GM-CSF and ipilimumab. This trial demonstrated significant benefit of the combination therapy over monotherapy (OS 17.5 months vs 12.7 months with an improved mortality hazard ratio of 0.64)^106^. For IL2, the trial results are more complicated as there are varied effects of the different multimeric combinations of the three components, IL2RA, IL2RB, and IL2RG on the balance between T effector and Treg actions. Despite encouraging phase II studies, the phase III study comparing nivolumab vs the combination of nivolumab and a pegylated interleukin-2 (IL-2) cytokine prodrug in advanced melanoma showed no benefit with the combination^107^.

Our results represent an unbiased genetic approach that again points to the GM-CSF and IL2/IL15 receptor systems as important to ICI response; but our genomic investigations may also shed further light as to the nuances of their biology. We showed that these blocking antibodies abrogated not only biomarkers of response (*CXCL9/SPP1* ratio and our 48 gene panel) but also the global transcriptomic effects of pharmacological aPD1 administration based on unbiased PCA analysis. Importantly, while aGM-CSF completely reversed the effects of aPD1 back to the untreated MC38 tumor configuration in the CCF1 (CC75×B6) responder genetic background, aIL2RB went further and completely reverted the transcriptional profile of the aPD1 treated tumors back to its baseline in the genetic non-responder (CC80×B6) host. This suggests that even in the presence of a TIME primed for response (potentially by IL2/IL15 pathways), certain cytokines like GM-CSF are necessary for the intratumor immune response induced by PD1 blockade. By contrast, other cytokines (such as the IL2R/IL15R system) may play a more foundational role, such as being indispensable for establishing a response propitious TIME. Our identification of *Il2rb*, the beta subunit of the multimeric IL2R system, and not the other IL2R components raises an intriguing possibility. Since the IL2RB can also associate with IL15RA and IL2RG in forming a highly functional heterotrimeric IL15 receptor complex, this suggests that the IL15 system may also be involved with our identified candidate which, if corroborated, could explain some of the variances in the clinical trials involving combination pegylated IL2 and ICIs. Indeed, the genes encoding the three subunits comprising both the high-affinity IL-2 receptor (*Il2ra, Il2rb, Il2rg*) and the IL-15 receptor (*Il15ra, Il2rb, Il2rg*) were co-expressed and significantly enriched in CTL clusters in responding MC38 tumors suggesting, at least, that the biochemical machinery is present for both signaling pathways in the major effector cells.

Interestingly, these results also suggest that different dosing regimens and more precise patient selection may improve outcomes. Because the primary effect of GM-CSF and IL2RB appears to be on the establishment of an ICI responsive TIME, it may be more efficacious to pretreat patients with the cytokines before ICI administration. The ECOG (E1608) trial gave GM-CSF for 14 days with aCTLA4 given on day 1. Our results suggest that there would be greater patient benefit if GM-CSF were given for the first 14 days and the ICI administered on the last day. When coupled with the 48 pan-QTL gene panel described herein, we would also expect the most advantageous outcomes to be seen in individuals with immune subtype 4 tumors but with lower scores with the 48 gene panel. By contrast, those with high gene panel scores would be likely be good responders to ICI and would see no benefit with adjunctive cytokine treatment. Moreover, given the molecular heterogeneity of the patient populations, any trial that did not stratify on biomarkers such as the *CXCL9/SPP1* ratio or our 48 pan-QTL gene panel could result in imbalances in the randomization that could incorrectly show that the GM-CSF and ICI combination had no benefit over monotherapy. This scenario is not merely hypothetical as the consistent efficacy of anti-EGFR therapeutics in non-small cell lung cancer was determined only after mutations in the kinase domain of EGFR were discovered as the critical predictive biomarker^108^.

Consistent with our finding of the importance of the MHC genes in defining the responder genotype, others have noted specific HLA haplotypes associated with ICI response in patients^27,29^. Furthermore, it has been observed in cancer patients treated with immune checkpoint inhibitors that HLA heterozygosity is associated with improved survival compared to homozygosity at any one of these loci. The explanation was that HLA allele heterozygosity enhances the ability of the immune system to recognize a wider range of antigens^27,109^. This was recently bolstered by large cohort studies of over 500,000 individuals from the UK Biobank and FinnGen resources that showed again HLA heterozygosity was significantly protective for the development of lung cancer^110^. However, it should also be considered that only particular heterozygous combinations of HLA haplotype may quantitatively expand anti-tumor immune responses. This has been demonstrated in mice where the heterozygous presence of a particular MHC can support the negative selection of T-cells in the thymus whose activity would otherwise be mediated by the opposing haplotype^111^.

Using single cell RNA analysis, the UK Biobank/FinnGen results also noted that smokers exhibited elevation of HLA-DRB1 in associated normal alveolar macrophages which can act as antigen presenting cells. When lung cancers were analyzed, they found that HLA II heterozygosity drove higher HLA-II expression. While the overexpression of certain MHC class I and class II genes (orthologs to HLA-A, HLA-DMA, HLA-DMB, HLA-DQA1, HLA-DQB1, HLA-DRA, HLA-DRB5, HLA-E, and HLA-F) were also associated with response in our CCF1 system, the impact of MHC heterozygosity appears to be more complicated. We found that homozygosity at the B6/129 H2^b^ MHC locus was associated with a nearly ten-fold increase in an immunodominant H2-K^b^-restricted MC38-specific CTL response than in non-responder strain tumors heterozygous at this locus. This states that with this model system, homozygosity at H2-K^b^ locus in both the germline and the tumor establishes a TIME with CTL responses that specifically target the MC38 tumor. The consequence of H2-K^b^ homozygosity in this situation is likely to be an experimental outlier based on the immunodominant nature of the MC38 CTL response. With these exceptional outliers (including the NOD strain that has a genetic antigen presenting cell defect) removed, the association between heterozygosity at the MHC locus and ICI response in our system is similar to the observations in humans of greater HLA heterozygosity being associated with favorable cancer outcomes.

Taken together, this work validates our CCF1/CCF1N1 system as a translationally relevant model system to identify host genes shaping the ICI-responsive TIME for any number of murine genetic models of a human cancer. The challenge with murine cancer models is that they are often not responsive to ICIs in the native inbred genetic background. However, we have found that all of the murine models we tested develop a range of responses. Thus, unlike other experimental systems, our approach identifies specific genetic backgrounds suitable for testing agents converting non-responders to responders, and vice versa, for any experimentally tractable murine adoptive transfer tumor model. Thus, with this CCF1/CCF1N1 system, we can genetically construct preclinical models that are “tuned” by targeted crosses for potentially any TIME composition over a range of expected responses. Such a platform will dramatically expand the experimental possibilities for discovering more precise immune checkpoint inhibitors and dosing protocols.

## Acknowledgements

This work was supported by an Aspire Award from The Mark Foundation for Cancer Research (Grant #19-036-ASP), the JAX Cancer Center’s Developmental Fund Award (P30 CA034196) and CATch Program, and the JAX Director’s Innovation Fund. The results published here are partially based upon data generated by the TCGA Research Network: https://www.cancer.gov/tcga. We gratefully acknowledge the contribution of the Single Cell Biology service, the Genome Technologies service, *In Vivo* Pharmacology section, the Histological Sciences service, and Cyberinfrastructure high performance computing resources at The Jackson Laboratory for expert assistance with the work described herein. These shared services are supported in part by the JAX Cancer Center (P30 CA034196). We gratefully acknowledge the provision of the MC38 cell line by Marcus Bosenburg (Yale University). We thank Ed Leiter, Susie Airhart, and Karolina Palucka for their thoughtful advice and guidance throughout the project, Nadia Rosenthal for manuscript review, and Anna Lisa Lucido and Jane Cha for assistance with graphics.

