## Supplementary Figures and Tables for "Mapping the genetic landscape establishing a tumor immune microenvironment favorable for anti-PD-1 response in mice and humans"

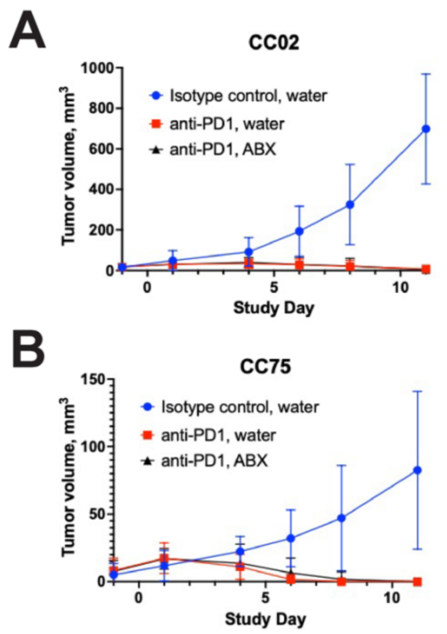


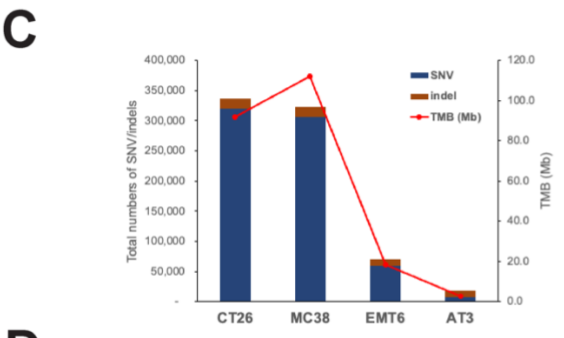


**D**

| | Deletion | Inversion | Tandem Duplication | Translocation | TDP status |
|---|---|---|---|---|---|
| **CT26** | 165 | 35 | 108 | 31 | TDP_group12mix |
| **MC38** | 93 | 32 | 68 | 15 | TDP_group2 |
| **EMT6** | 82 | 27 | 74 | 12 | TDP_group2 |
| **AT3** | 37 | 12 | 7 | 19 | non TDP |

**Figure S1.** (A-B) Responder strain (CC02 and CC075 F1) mice given broad spectrum antibiotics (vancomycin, streptomycin, ampicillin, and colistin) *ad libitum* in drinking water display no loss in response to PD1 blockade. (C) SNP/indel count and tumor mutation burden (per megabase) in each tumor model. (D) Summary of single base and structural variations, and TDP assessment, quantified in each tumor model.

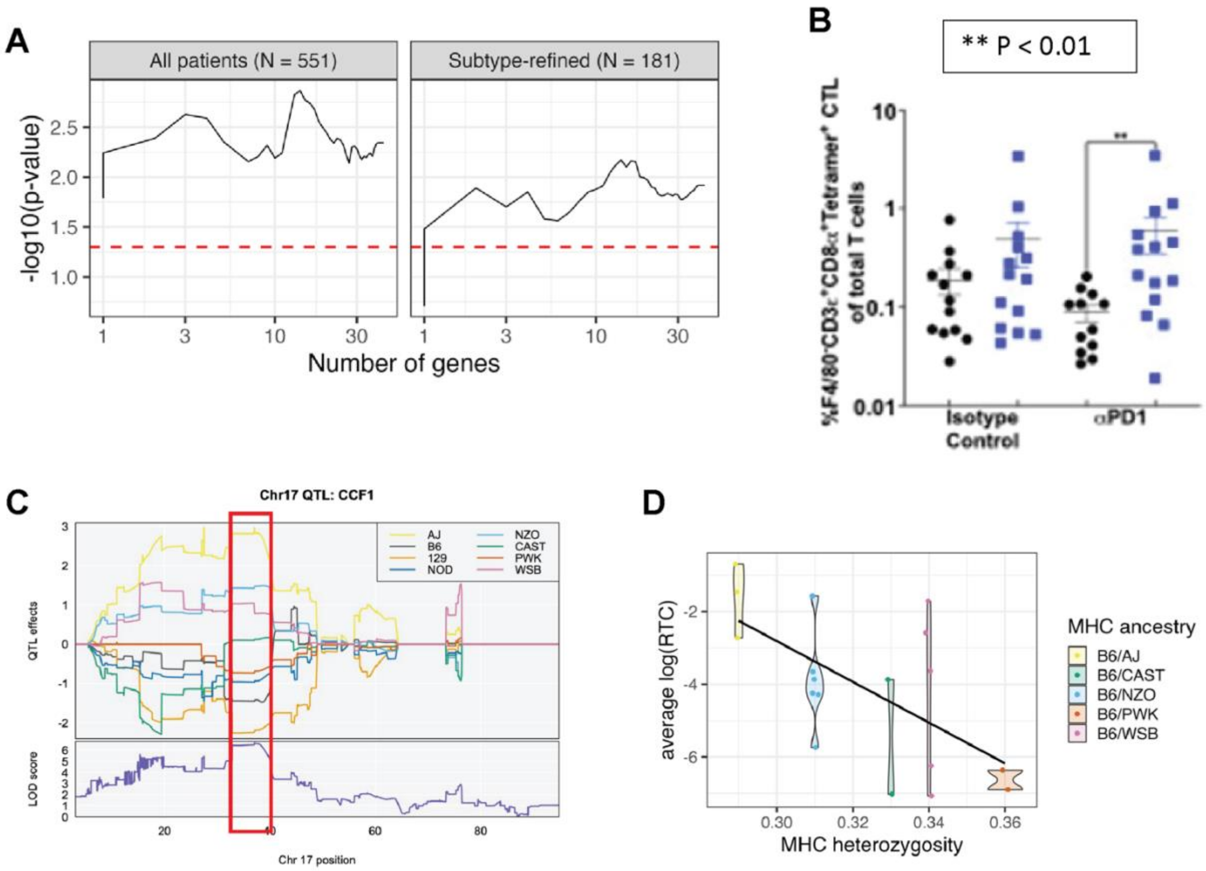


**Figure S2.** (A) Statistical significance levels of different numbers of genes in ICI response prognostic gene set for all patients (left) or subtype 4-restricted patients (right). (B) FACS data showing MC38-specific CTL as a percentage of total T cells. black circles indicate measurements taken from non-responder strain mice (CC36 F1, CC79 F1, and CC80 F1), blue boxes indicate measurements taken from responder strain mice (CC01 F1, CC02 F1, and CC075 F1). (C) Top panel shows QTL effects plots showing the effect on RTC of carrying a haplotype derived from each of the eight founder parental lines of the CC. Lower number indicates a lower value of RTC, which is associated with better response. Bottom panel shows LOD score across the locus from QTL mapping, as in Figure 2. Red box indicates approximate boundaries of the MHC locus. (D) Plot of ICI response as measured by RTC, stratified by each CCF1 line’s ancestry at the MHC locus, versus MHC heterozygosity. As in (C), a lower RTC number is associated with better response.

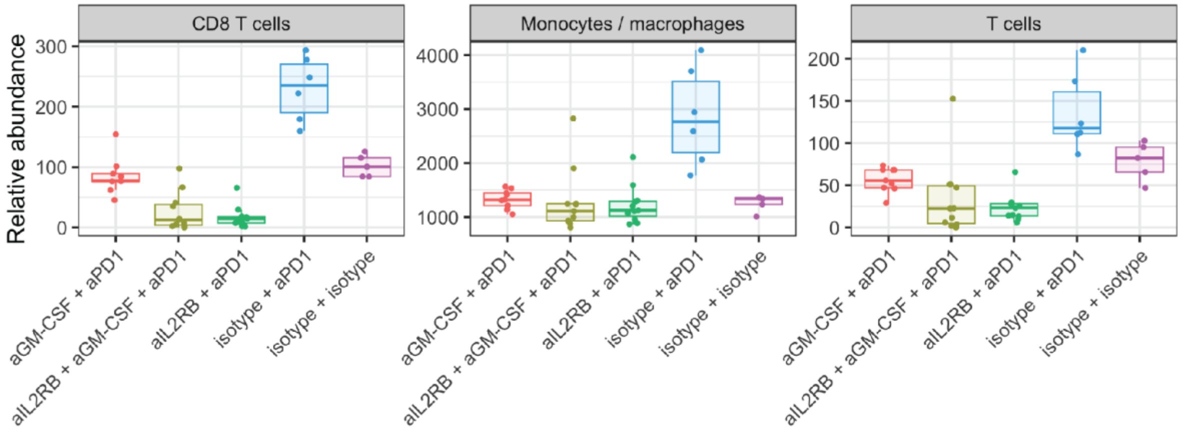


**Figure S3.** Bulk RNA-Seq deconvolution using mMCP-Counter on samples of MC38 tumors from CC75F1 mice treated with blocking antibodies to PD1, GM-CSF, and/or IL2RB.

| JAX strain number | Strain name |
|---|---|
| 664 | C57BL/6J |
| 651 | BALB/cJ |
| 21238 | CC001 |
| 21236 | CC002 |
| 21237 | CC003 |
| 20944 | CC004 |
| 20945 | CC005 |
| 29625 | CC007 |
| 26971 | CC008 |
| 18856 | CC009 |
| 21889 | CC010 |
| 18854 | CC011 |
| 28409 | CC012 |
| 18859 | CC015 |
| 24684 | CC016 |
| 22870 | CC017 |
| 25131 | CC023 |
| 21891 | CC024 |
| 18857 | CC025 |
| 25130 | CC027 |
| 20946 | CC032 |
| 25910 | CC033 |
| 25127 | CC036 |
| 26426 | CC044 |
| 24683 | CC057 |
| 27298 | CC058 |
| 26427 | CC060 |
| 23830 | CC065 |
| 25908 | CC068 |
| 18855 | CC074 |
| 27293 | CC075 |
| 25989 | CC078 |
| 25990 | IL6411(CC079) |
| 25988 | IL6573(CC080) |

**Table S1: Mouse strains used in this study including links for sourcing mice.**

| | | Non-responders, αPD1 versus ISO | | | | Responders, aPD1 versus ISO | | | | Responders versus Non-responders (ISO) | | Responders versus Non-responders (αPD1) | |
|---|---|---|---|---|---|---|---|---|---|---|---|---|---|
| Population or cluster | FACS or scRNAseq cluster | ISO | anti-PD1 | Fold change αPD1:ISO | Significant difference? | ISO | anti-PD1 | Fold change αPD1:ISO | Significant difference? | Fold Change R:NR | Significant difference RvsNR? | Fold Change R:NR | Significant difference RvsNR? |
| %CD45$^{+}$ of Live Cells | FACS | 66.2+/-3.15 | 72.85+/-2.89 | 1.1 | No | 68.16+/-2.82 | 64.69+/-4.41 | 0.95 | No | 1.03 | No | 0.89 | No |
| %macrophages of CD45$^{+}$ | FACS | 70.38+/-3.03 | 71.03+/-2.11 | 1.01 | No | 68.49+/-2.44 | 69.08+/-2.63 | 1.01 | No | 0.97 | No | 0.97 | No |
| %T cells of CD45$^{+}$ | FACS | 19.75+/-2.12 | 21.39+/-2.87 | 1.08 | No | 20.57+/-2.28 | 20.11+/-2.47 | 0.98 | No | 1.04 | No | 0.94 | No |
| %CD4$^{+}$ T cells of total T cells | FACS | 3.39+/-0.79 | 3.22+/-0.62 | 0.95 | No | 2.15+/-0.30 | 2.02+/-0.20 | 0.94 | No | 0.63 | No | 0.63 | No |
| %CTLs of total T cells | FACS | 2.16+/-0.43 | 1.57+/-0.36 | 0.73 | No | 2.82+/-0.55 | 3.37+/-0.74 | 1.2 | No | 1.31 | No | 2.15 | Yes, P*=0.0270 |
| %Tetramer$^{+}$ CTLs of total T cells | FACS | 0.19+/-0.06 | 0.09+/-0.02 | 0.47 | No | 0.48+/-0.23 | 0.58+/-0.24 | 1.21 | No | 2.53 | No | 6.44 | Yes, P**=0.0037 |
| %Tetramer$^{+}$PD1$^{+}$ CTLs of total T cells | FACS | 0.06+/-0.02 | 0.01+/-0.003 | 0.17 | Yes P***=0.0006 | 0.31+/-0.18 | 0.044+/-0.01 | 0.14 | No | 5.17 | No | 3.38 | Yes, P*=0.0108 |
| %CD206$^{-}$PD-L1$^{high}$MHCII$^{high}$ macrophages of total macrophages | FACS | 12.80+/-1.41 | 17.41+/-3.15 | 1.36 | No | 17.66+/-1.94 | 26.09+/-4.61 | 1.48 | No | 1.38 | Yes, P*=0.0427 | 1.5 | No |
| %CD206$^{+}$ macrophages of total macrophages | FACS | 31.30+/-3.57 | 23.39+/-3.46 | 0.75 | No | 31.63+/-4.60 | 27.65+/-4.41 | 0.87 | No | 1.01 | No | 1.18 | No |
| %Exhausted CTL of total dataset | 3 | 2.29+/-1.04 | 0.67+/-0.2 | 0.29 | No | 5.041+/-1.17 | 4.81+/-1.2 | 0.95 | No | 2.2 | Yes, P*=0.0435 | 7.18 | Yes, P****<0.0001 |
| %Ifng$^{+}$ CTL of total dataset | 13 | 0.31+/-0.08 | 0.23+/-0.057 | 0.74 | No | 1.6+/-0.66 | 0.73+/-0.10 | 0.46 | No | 5.16 | Yes, P*=0.0399 | 3.17 | Yes, P**=0.0021 |
| %IFNγ-stimulated macrophages of total dataset | 2, 5, 26 | 3.61+/-0.88 | 3.38+/-0.46 | 0.94 | No | 8.26+/-1.15 | 17.22+/-2.85 | 2.08 | Yes P**=0.0029 | 2.29 | Yes, P**=0.0057 | 5.09 | Yes, P****<0.0001 |
| %Cd209a$^{+}$ DC of total dataset | 14 | 6.68+/-0.59 | 5.65+/-0.56 | 0.85 | No | 4.44+/-0.62 | 4.21+/-0.58 | 0.95 | No | 0.66 | Yes, P*=0.0172 | 0.75 | No |
| %cDC1 of total dataset | 11 | 1.78+/-0.33 | 1.61+/-0.21 | 0.9 | No | 1.395+/-0.24 | 1.68+/-0.2 | 1.2 | No | 0.78 | No | 1.04 | No |
| %Xcr1$^{+}$Clec9a$^{+}$ DC of total dataset | 6 | 1.59+/-0.3 | 1.31+/-0.24 | 0.82 | No | 1.21+/-0.22 | 1.34+/-0.2 | 1.11 | No | 0.76 | No | 1.03 | No |
| %plasmacytoid DC of total dataset | 18 | 0.56+/-0.12 | 0.4+/-0.1 | 0.71 | No | 0.51+/-0.15 | 0.56+/-0.19 | 1.1 | No | 0.91 | No | 1.4 | No |

**Table S2: Overview of flow cytometry results.**

| mouse chromosome | mouse gene symbol |
|---|---|
| 5 | ***Acox3***, ***Add1***, ***Afap1***, *Emilin1*, ***Fndc4***, ***Gm1673***, ***Ift172***, ***Mpv17***, ***Mxd4***, ***Nrbp1***, ***Sh3bp2***, ***Stk32b***, ***Ywhah*** |
| 9 | ***AB124611***, ***Acp5***, *Anln*, ***Cadm1***, *Cbl*, ***Cd3d***, ***Cd3e***, ***Cd3g***, *Cdon*, *Dync2h1*, *Fat3*, ***Fli1***, ***Icam1***, *Ift46*, ***Il10ra***, *Naalad2*, *Ncam1*, ***Pknox2***, *Prkcsh*, *Rexo2*, ***S1pr2***, *Slc37a2*, *Slc37a4*, ***St14***, *Tagln*, *Thy1*, *Tyk2*, *Ubash3b*, *Zfp426* |
| 15 | ***1700088E04Rik***, ***Apobec3***, ***Apol6***, *C1qtnf6*, ***Csf2rb***, ***Csf2rb2***, ***Cyth4***, *Elfn2*, ***Fam83f***, *Gpaa1*, ***Grap2***, ***Gtpbp1***, ***Il2rb***, ***Kdelr3***, *Pdgfb*, ***Rac2***, ***Sh3bp1***, *Syngr1* |
| 17 | *Abca3*, *Abcg1*, *Aif1*, *Atp6v1g2*, *C2*, *Ddah2*, *Ddr1*, ***Fgd2***, *Fkbp5*, *Fpr2*, ***Gm11127***, *Gm8909*, *Gpsm3*, ***H2-Aa***, *H2-Ab1*, *H2-D1*, *H2-DMa*, ***H2-DMb1***, ***H2-Ea***, *H2-Eb1*, *H2-K1*, *H2-Q7*, ***H2-T23***, *Lmf1*, *Lsm2*, *Lst1*, ***Ltb***, *Mmp25*, *Msln*, *Myo1f*, *Nme4*, *Pglyrp2*, *Psmb8*, ***Psmb9***, ***Rasal3***, *Rps18*, *Sik1*, *Syngap1*, ***Tap1***, *Tap2*, *Tead3*, *Tnf*, ***Ubash3a***, ***Ubd*** |

**Table S3:** Mouse orthologs of the human genes prioritized using our CST algorithm predicted to have a major role establishing a tumor immune microenvironment favorable for aPD1 response in mice and humans
